# Atomic structures of respiratory complex III_2_, complex IV and supercomplex III_2_-IV from vascular plants

**DOI:** 10.1101/2020.08.30.274431

**Authors:** María Maldonado, Fei Guo, James A. Letts

## Abstract

Mitochondrial complex III (CIII_2_) and complex IV (CIV), which can associate into a higher-order supercomplex (SC III_2_+IV), play key roles in respiration. However, structures of these plant complexes remain unknown. We present atomic models of CIII_2_, CIV and SC III_2_+IV from *Vigna radiata* determined by single-particle cryoEM. The structures reveal plant-specific differences in the MPP domain of CIII_2_ and define the subunit composition of CIV. Conformational heterogeneity analysis of CIII_2_ revealed long-range, coordinated movements across the complex, as well as the motion of CIII_2_’s iron-sulfur head domain. The CIV structure suggests that, in plants, proton translocation does not occur via the H-channel. The supercomplex interface differs significantly from that in yeast and bacteria in its interacting subunits, angle of approach and limited interactions in the mitochondrial matrix. These structures challenge long-standing assumptions about the plant complexes, generate new mechanistic hypotheses and allow for the generation of more selective agricultural inhibitors.

## Introduction

The canonical mitochondrial electron transport chain (mETC), composed of four integral membrane protein complexes (complexes I-IV; CI-CIV) located in the inner mitochondrial membrane (IMM), transfers electrons from NADH and succinate to molecular oxygen. The concomitant pumping of protons (H^+^) across IMM the establishes an electrochemical proton gradient that is used by ATP synthase to produce ATP^[1]^. Whereas the atomic details of the respiratory complexes and several supercomplexes (higher-order complex assemblies) are known for yeast, mammals and bacteria^[2–13]^, the high-resolution structural details of the respiratory complexes and supercomplexes of plants have remained mostly unknown.

Complex III (CIII_2_), also called the cytochrome *bc*_1_ complex or ubiquinol-cytochrome *c* oxidoreductase, is an obligate dimer that transfers electrons from ubiquinol in the IMM (reduced by CI, CII or alternative NADH dehydrogenases) to soluble cytochrome *c* in the IMS^[1]^. This redox reaction is coupled to the pumping four H^+^ to the IMS. CIII_2_ is composed of three core subunits present in all organisms (cytochrome *b*, COB; cytochrome *c*_1_, CYC1; and the iron-sulfur “Rieske” subunit, QCR1), as well as a varying number of accessory subunits present in eukaryotes^[14–16]^. Each CIII monomer contains one low-potential heme *b* (*b*_L_) and one high-potential heme *b* (*b*_H_) in COB, a heme *c* in CYC1, an 2Fe-2S iron-sulfur cluster in QCR1, as well as two quinone-binding sites (Q_P_ and Q_N_ close to the positive/IMS and negative/matrix sides respectively) in COB. Given that CIII_2_ is a dimer, these sites in each CIII monomer are symmetrical within the dimer in isolation. However, the symmetry may be broken when CIII assembles into asymmetrical supercomplexes^[5, 11, 17]^. CIII_2_’s redox and proton pumping occur via the “Q cycle” mechanism^[18]^, which allows for efficient electron transfer between ubiquinol (a two-electron donor) and cyt *c* (a one-electron acceptor). To this end, one electron is transferred from ubiquinol in the Q_p_ site to the 2Fe-2S in the QCR1 head domain. The head domain then undergoes a large conformational swinging motion from its “proximal” position close to the Q_P_ site to a “distal” CYC1 binding site adjacent to heme *c*_1_^[19]^. The electron is then transferred *via* heme *c_1_* to cyt *c* bound to CIII_2_ in the IMS. Of note, the QCR1 head domain belongs to the opposite CIII protomer relative to COB and CYC1. The second electron donated by ubiquinol is transferred *via* hemes *b*_L_ and *b*_H_ to a quinone in the Q_N_ site, reducing it to semi-ubiquinone. The cycle is repeated to regenerate ubiquinol in the Q_N_ site, ultimately reducing two molecules of cyt *c* and pumping four protons.

In eukaryotes, the large CIII_2_ accessory subunits exposed to the mitochondrial matrix have homology to mitochondrial processing peptidases (MPP) of the pitrilysin family^[20]^. These metalloendopeptidases –composed of an active *β* subunit and an essential but catalytically inactive *α* subunit –cleave the mitochondrial signal sequences of proteins that are imported into the mitochondria^[20]^. Whereas in yeast the CIII_2_ accessory subunits with MPP homology (ScCor1/Cor2) have completely lost MPP enzymatic activity, the mammalian CIII_2_ homologue (UQCR1/UQCR2 heterodimer) retains basal activity to only one known substrate (the Rieske subunit)^[20, 21]^. Hence, in yeast and mammals this enzymatic activity is carried out by soluble MPP heterodimers in the mitochondrial matrix. In contrast, in vascular plants, there is no additional soluble MPP enzyme, and all MPP activity is provided by the CIII_2_ MPP heterodimer (MPP-*α*/*β*). Thus, in plants CIII_2_ serves a dual role as a respiratory enzyme and a peptidase^[22–26]^. The reasons for and bioenergetic implications of this dual function remain unknown.

Complex IV (CIV), also called cytochrome *c* oxidase, transfers electrons from cyt *c* to molecular oxygen, reducing it to water^[1]^. The redox reaction is coupled to the pumping of four protons into the IMS. Like CIII_2_, CIV is composed of three core subunits (COX1, COX2, COX3) and a variable number of accessory subunits, depending on the organism. Electrons are transferred from cyt *c* to oxygen via Cox1’s dinuclear copper (Cu_A_), heme *a* and copper-associated heme *a*_3_ (Cu_B_, binuclear center). The passage of protons from the matrix to the IMS occurs through distinct “channels” formed by protonatable amino acid residues^[27–30]^. It is currently believed that, whereas yeast CIV pumps protons through the K and D transfer pathways (named after key amino acid residues in each pathway), mammalian CIV uses an H-channel in addition to the K and D channels^[27–31]^. The contribution of the K, D, H pathways in plant CIV has not been characterized.

In the IMM, respiratory complexes can be found as separate entities or as higher-order assemblies known as supercomplexes^[32]^. Supercomplexes of various stoichiometries between CIII_2_ and CIV (e.g. CIII_2_-CIV, CIII_2_-CIV_2_) have been seen^[32–34]^. In plants, the CIII_2_-CIV supercomplex of highest abundance is a single CIII dimer with a single CIV monomer (SC III_2_+IV)^[33]^. High-resolution structures of the model yeast *Saccharomyces cerevisiae* and *Mycobacterium smegmatis* CIII_2_-CIV supercomplex (SC III_2_+IV_2_) have recently been determined^[35–38]^. Although there is currently no high-resolution structure for a mammalian CIII_2_-CIV supercomplex, the supercomplex between CI, CIII_2_ and CIV (SC I+III_2_+IV, the respirasome) shows a distinct interaction interface between CIII_2_ and CIV relative the SC III_2_+IV_2_ from yeast and bacteria^[4, 5]^. Similar to that seen in comparative tomographic studies of SC I+III_2_+IV^[39]^, the above SC III_2_+IV studies revealed that, while the general configuration of the individual CIII_2_ and CIV are conserved, the location of the binding interface between CIII_2_ and CIV in the supercomplex is divergent, with different subunits involved in the different organisms. Structures for CIII_2_, CIV or their supercomplex are not currently available for the plant kingdom.

Although it was initially hypothesized that supercomplexes would allow for direct substrate channeling between complexes, evidence has mounted against this view^[4, 5, 11, 40–43]^. Instead, supercomplexes may have roles improving the stability of the complexes, providing kinetic advantages to the electron transfer or reducing the production of reactive oxygen species or of aggregates in the IMM^[17, 44]^.

Here we present the cryoEM structures of CIII_2_ and SC III_2_+IV from the vascular plant *Vigna radiata* (mung bean) at nominal resolutions of 3.2 Å and 3.8 Å, respectively. Using focused refinements around CIV, we achieved a nominal resolution for CIV of 3.8 Å. The structures reveal plant CIII_2_ and CIV’s active sites, as well as the plant-specific configuration of several CIII_2_ and CIV subunits. The structures also show the SC III_2_+IV binding interface and orientation, which is unique to plants. Additionally, using cryoEM 3D-conformation variability analysis^[45]^, we were able to visualize the swinging motion of CIII_2_’s QCR1 head domain at 5 Å resolution in the absence of substrate or inhibitors. We also observed complex-wide coupled conformational changes in the rest of the CIII_2_. These results question long-standing assumptions, generate new mechanistic hypotheses and allow for the development of more selective agricultural inhibitors of plant CIII_2_ and CIV.

## Results

Mitochondria were isolated from etiolated *V. radiata* hypocotyls. The electron transport chain complexes were extracted using the gentle detergent digitonin. The extracted complexes were further stabilized the with amphipathic polymers (amphipol) and separated on a sucrose gradient. Given our interest in respiratory complex I^[46]^, we pooled fractions containing NADH-dehydrogenase activity and set up cryoEM grids. Upon 2D classification of the particles in the micrographs, and supported by mass spectrometry data (Table 1), it became evident that the pooled fractions contained not only a CI intermediate^[46]^, but also CIII_2_ and SC III_2_+IV. Therefore, CIII_2_ and SCIII+CIV were purified *in silico*, in an example of “bottom-up structural proteomics” ^[47]^ on a partially purified sample. Processing of the CIII_2_ and SCIII_2_+IV particles resulted in near-atomic reconstructions of CIII_2_ at 3.2 Å, of CIV at 3.8 Å and of SCIII_2_+IV at 3.8 Å (Figure 1, Videos 1-3, Figure 1-Figure Supplements 1-3, Tables. 2-6).

**Figure 1.**
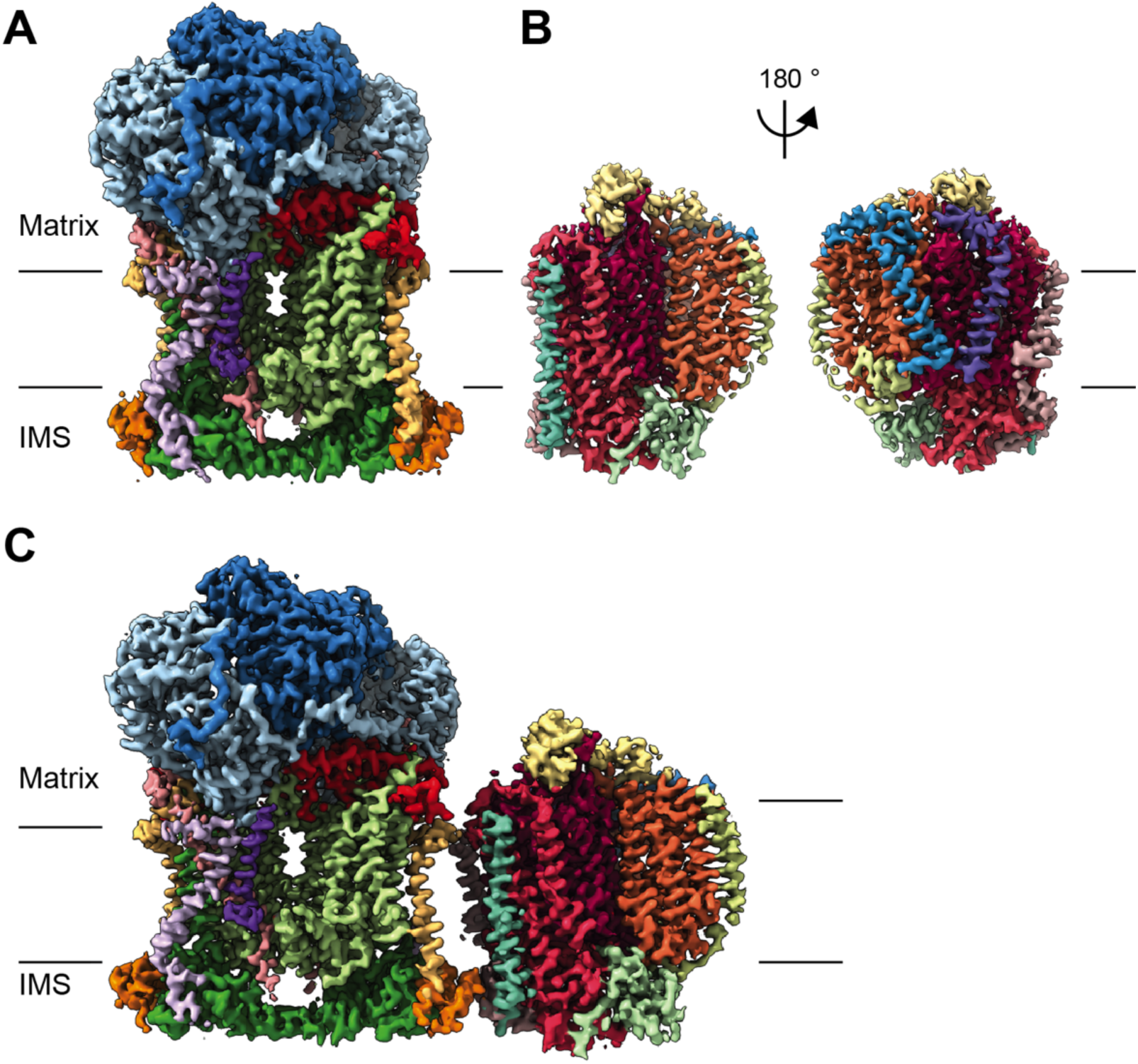
CryoEM reconstructions for *V. radiata* mitochondrial CIII_2_, CIV and SCIII_2_+IV. (A) CryoEM density map for CIII_2_ in isolation (not assembled into a supercomplex; see also Figure 1-Figure Supplement 1 and Video 1). (B) Density map for CIV, obtained from re-centered focused refinements of CIV in the supercomplex (see also Figure 1- Figure Supplement 2 and Video 2). (C) Composite map of SC III_2_+IV, assembled by combining CIII_2_ and CIV focused refinements from the SC particles (see also Figure 1-Figure Supplements 1 and 2 and Video 3). Volume surfaces are colored by subunit (see also Videos 1-3 and Figures 2 and 5). The approximate position of the matrix and IMS sides of the membrane are shown with black lines.

**Table 1.**
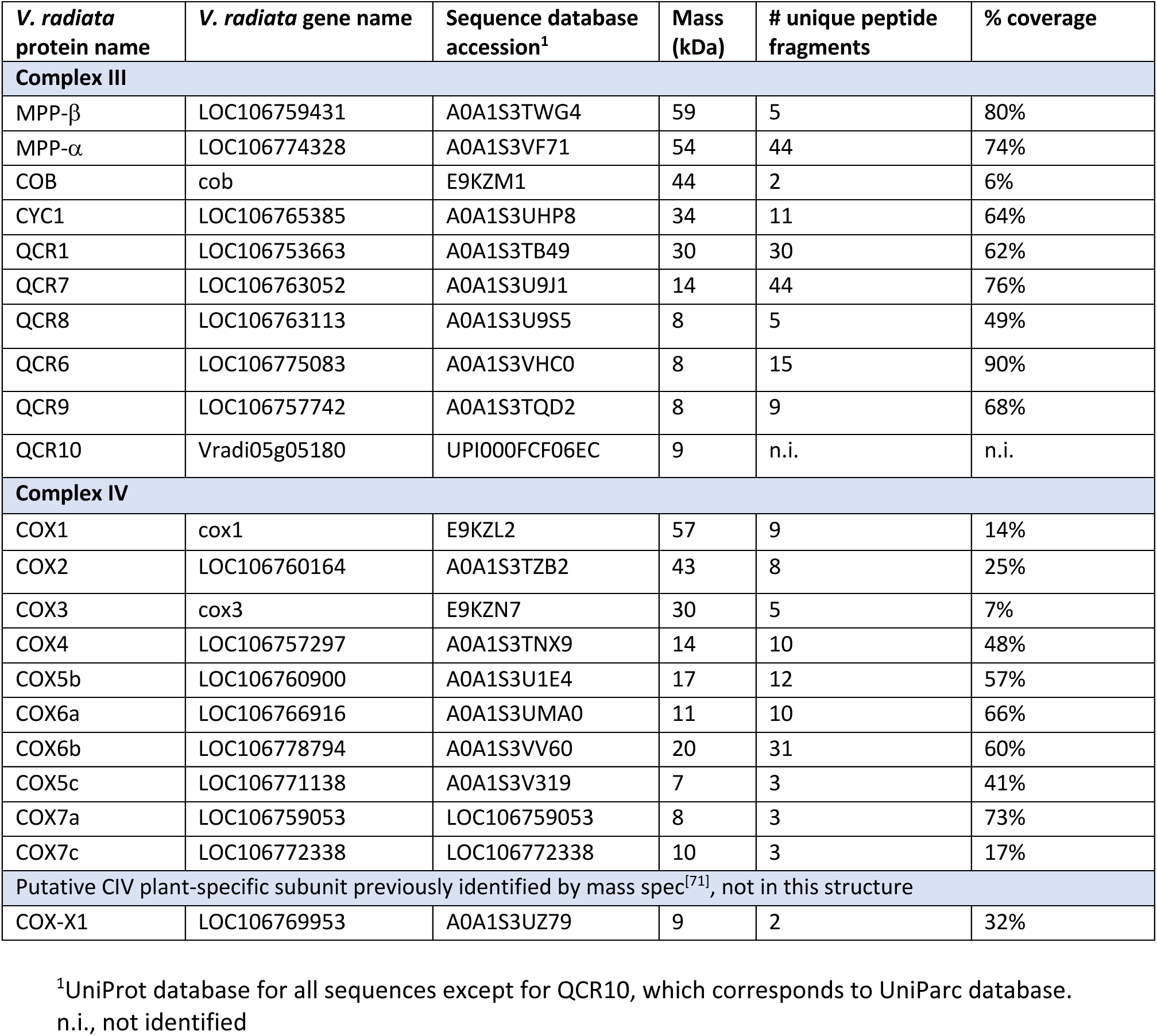
Mass spectrometry identification of V. radiata CIII and CIV subunits.

**Table 2.**
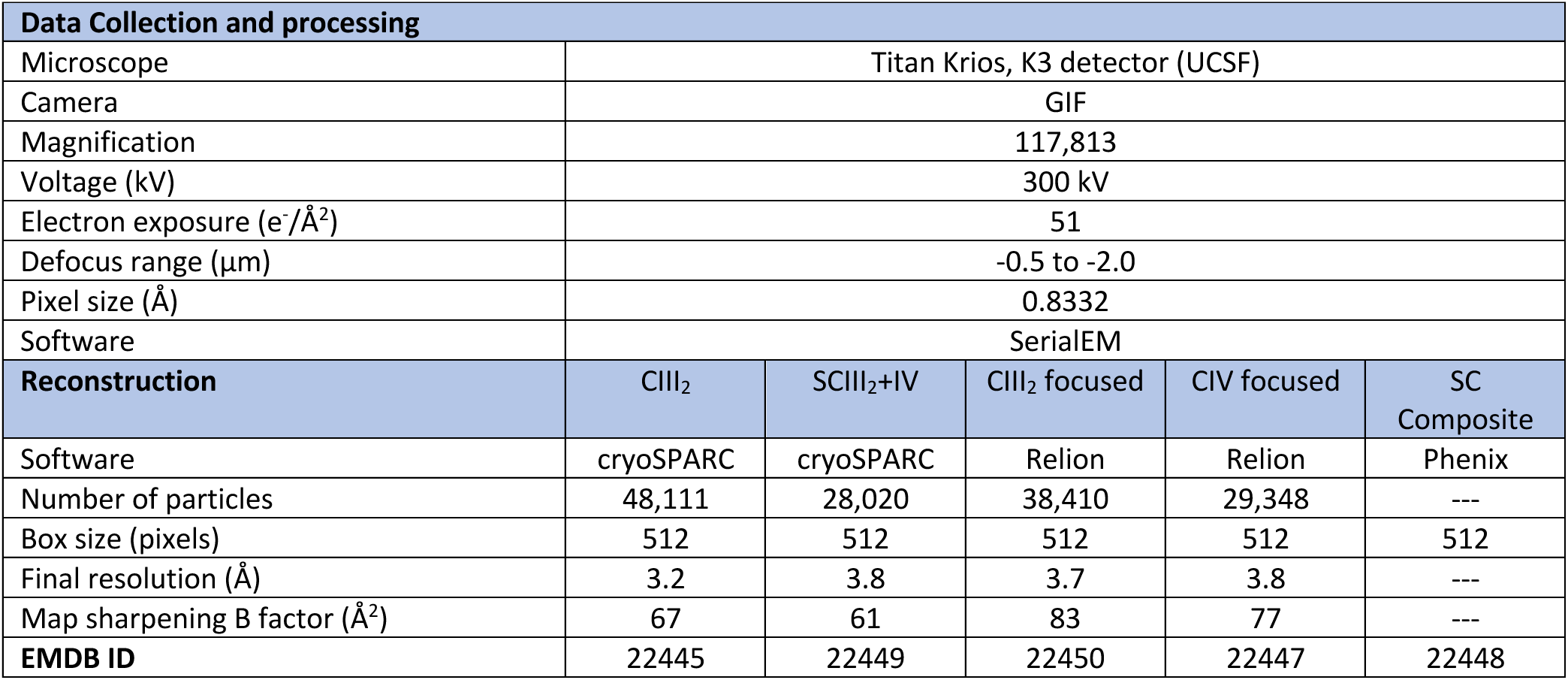
Cryo-EM data collection and reconstruction statistics.

| Data Collection and processing |  |  |  |  |  |
| --- | --- | --- | --- | --- | --- |
| Microscope | Titan Krios, K3 detector (UCSF) |  |  |  |  |
| Camera | GIF |  |  |  |  |
| Magnification | 117,813 |  |  |  |  |
| Voltage (kV) | 300 kV |  |  |  |  |
| Electron exposure (e<sup>-</sup>/Å<sup>2</sup>) | 51 |  |  |  |  |
| Defocus range (μm) | -0.5 to -2.0 |  |  |  |  |
| Pixel size (Å) | 0.8332 |  |  |  |  |
| Software | SerialEM |  |  |  |  |
| Reconstruction | CIII<sub>2</sub> | SCIII<sub>2</sub>+IV | CIII<sub>2</sub> focused | CIV focused | SC Composite |
| Software | cryoSPARC | cryoSPARC | Relion | Relion | Phenix |
| Number of particles | 48,111 | 28,020 | 38,410 | 29,348 | --- |
| Box size (pixels) | 512 | 512 | 512 | 512 | 512 |
| Final resolution (Å) | 3.2 | 3.8 | 3.7 | 3.8 | --- |
| Map sharpening B factor (Å<sup>2</sup>) | 67 | 61 | 83 | 77 | --- |
| **EMDB ID** | 22445 | 22449 | 22450 | 22447 | 22448 |

**Table 3.**
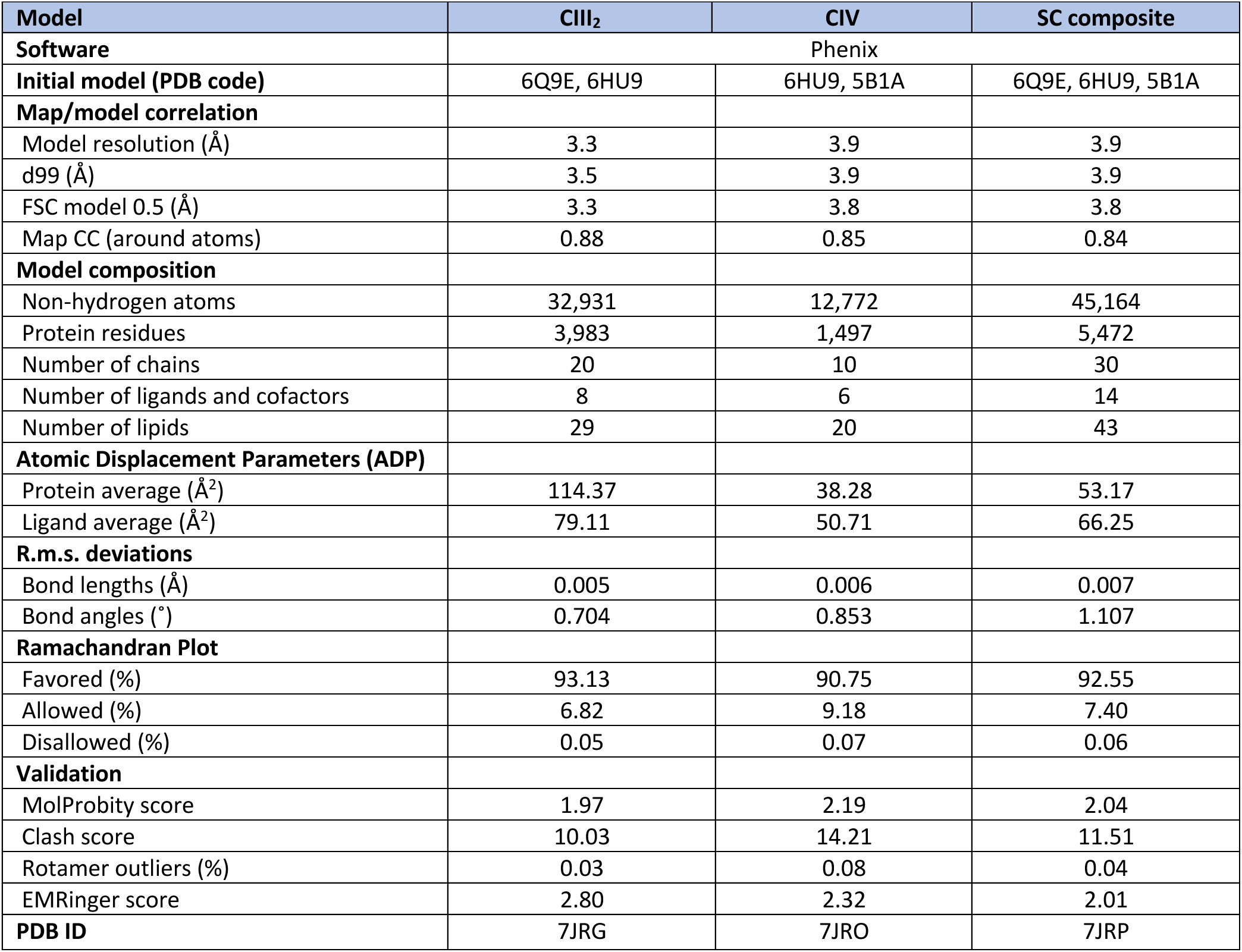
Model refinement and validation.

### Complex III dimer (CIII2)

#### Overall structure and ligands

The *V. radiata* (Vr) structure confirms that VrCIII_2_ contains 10 subunits per CIII monomer (3 core subunits and 7 accessory subunits), similar to yeast and mammals (Figure 2, Figure 2- Figure Supplement 1). As in mammals and yeast, VrCIII_2_ contains 13 transmembrane helices per protomer: 8 from COB and 1 from each of CYC1 (cyt *c*_1_), QCR1, QCR8, QCR9 and QCR10 (Table 5). The MPP domain, composed of two MPP-*α*/*β* heterodimers (one per CIII protomer), extends into the matrix. The C-terminus of CYC1 (six *α*-helices and a 2-strand *β*-sheet), the entire QCR6 subunit and QCR1’s head domain extend into the intermembrane space (IMS). Consistent with their known flexibility and conformational heterogeneity, the linker and head domains of QCR1 (Rieske iron-sulfur subunit) were disordered in our reconstruction and an atomic model was not produced for this region of QCR1. (Throughout the manuscript, we use plant subunit nomenclature unless otherwise stated; see Table 4 for details and name equivalence in other organisms.)

**Figure 2.**
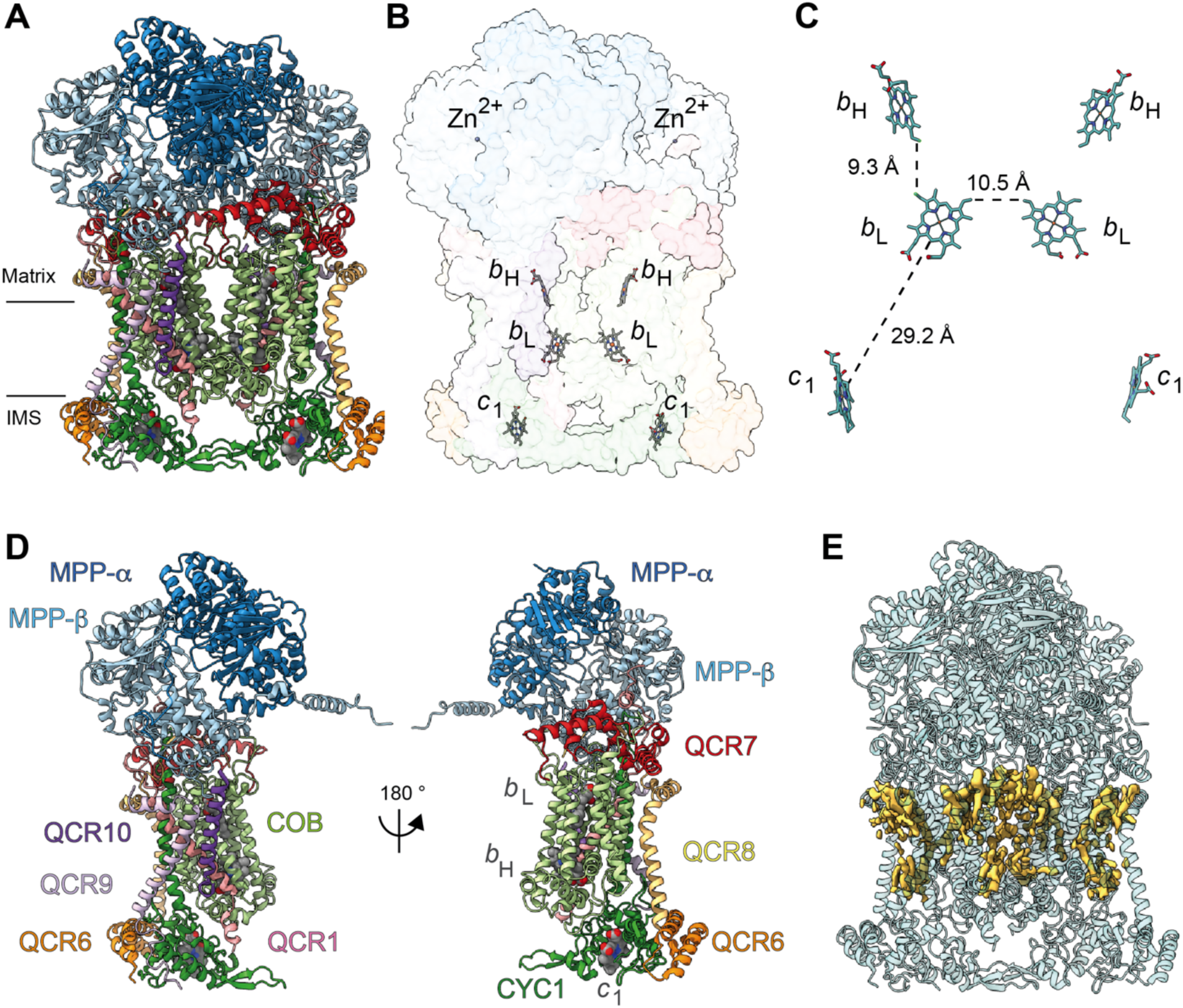
Overview of plant CIII_2_ atomic model. (A) CIII_2_ in cartoon representation with co-factors in sphere representation. The approximate position of the inner mitochondrial membrane is shown with black lines, and the matrix and inter-membrane space (IMS) sides are labelled. (B) Position of the observed CIII_2_ co-factors. Note that the iron-sulfur groups are not shown because the flexible head domain of the iron-sulfur protein is disordered in our cryoEM density. CIII_2_ shown in transparent surface representation, cofactors in stick representation. (C) Distances between the heme groups are shown, calculated edge-to-edge to the macrocyclic conjugated system. (D) Each CIII and co-factor are shown as in (A) with subunits labeled. The CIII monomers are separated for clarity and the two-fold symmetry axis is indicated. (E) Density consistent with lipids (yellow) is shown overlaid on the CIII_2_ cartoon model (transparent teal). *b*_H_, high potential heme *b*; *b*_L_, low potential heme *b*; *c*_1_, heme *c*_1_; IMS, intermembrane space.

**Table 4.** CIII and CIV subunit homologues in plants, yeast and mammals. . *V. radiata* homologues were obtained by performing BLASTp searches of the *Arabidopsis thaliana* genes^[84]^. Mammalian and yeast homologues were obtained from ref. ^[35]^,^[85]^. Additional BLASTp searches were performed as needed.

| Chain ID<sup>1</sup> | Subunit name<sup>2</sup> |  |  |  | Gene<sup>2</sup> |  |  |  |
| --- | --- | --- | --- | --- | --- | --- | --- | --- |
|  | *V. radiata* | *A. thaliana* | *S. cerevisiae* | *B. taurus* | *V. radiata*<sup>3</sup> | *A. thaliana* | *S. cerevisiae* | *B. taurus* |
| **Complex III** |  |  |  |  |  |  |  |  |
| A,M | MPP-β | MPP-b | Cor1 | UQCRC1 | LOC106759431 | At3g02090 | *COR1* | *UQCRC1* |
| B,N | MPP-α | MPP-a | Cor2 | UQCRC2 | LOC106774328 | At1g51980 | *QCR2* | *UQCRC2* |
| C,O | COB | COB | Cytb | MT-CYB | cob | AtMg00220 | *COB* | *MT-CYB* |
| D,P | CYC1 | CYC1 | Cytc1 | CYC1 | LOC106765385 | At5g40810 | *CYT1* | *CYC1* |
| E,Q | QCR1 | UCR1 | Rip1 | UQCRFS1 | LOC106753663 | At5g13430 | *RIP1* | *UQCRFS1* |
| F,R | QCR7 | QCR7 | Qcr7 | UQCRB | LOC106763052 | At4g32470 | *QCR7* | *UQCRB* |
| G,S | QCR8 | UCRQ | Qcr8 | UQCRQ | LOC106763113 | At3g10860 | *QCR8* | *UQCRQ* |
| H,T | QCR6 | QCR6 | Qcr6 | UQCRH | LOC106775083 | At1g15120 | *QCR6* | *UQCRH* |
| J,V | QCR9 | QC9 | Qcr9 | UQCR10 | LOC106757742 | At3g52730 | *QCR9* | *UQCR10* |
| K,W | QCR10 | UCRY | Qcr10 | UQCR11 | Vradi05g05180 | At2g40765 | *QCR10* | *UQCR11* |
| **Complex IV** |  |  |  |  |  |  |  |  |
| a | COX1 | COX1 | Cox1 | COX1 | cox1 | AtMg01360 | *COX1* | *COX1* |
| b | COX2 | COX2 | Cox2 | COX2 | LOC106760164 | AtMg00160 | *COX2* | *COX2* |
| c | COX3 | COX3 | Cox3 | COX3 | cox3 | AtMg00730 | *COX3* | *MT-CO3* |
| d | COX4 | COX-X2 | Cox5a | COX4 | LOC106757297 | At4g00860 | *COX5a* | *CO4IL1* |
| e | COX5b | COX5b | Cox4 | COX5B | LOC106760900 | At3g15640 | *COX4* | *COX5B* |
| f | COX6a | COX6a | Cox13 | COX6A | LOC106766916 | At4g37830 | *COX13* | *COX6A1* |
| g | COX6b | COX6b | Cox12 | COX6B | LOC106778794 | At1g22450 | *COX12* | *COX6B1* |
| h | COX5c | COX5c | Cox9 | COX6C | LOC106771138 | At2g47380 | *COX9* | *COX6C1* |
| i | COX7a | COX-X4 | Cox7 | COX7A | LOC106759053 | At4g21105 | *COX7* | *CO7A1* |
| j | COX7c | COX-X3 | Cox8 | COX7C | LOC106772338 | At1g72020 | *COX8* | *COX7C* |
| - | - | - | Cox6 | COX5A | - | - | *COX6* | *COX5A* |
| - | - | - | Cox26 | - | - | - | *COX26* | - |
| - | - | - | - | COX7B | - | - | - | *COX7B* |
| - | - | - | - | COX8B | - | - | - | *COX8B* |
| Putative CIV plant-specific subunits identified in previous mass spec studies<sup>[71, 86]</sup> not seen in our structure |  |  |  |  |  |  |  |  |
| *A. thaliana* gene | *A. thaliana* protein |  |  | *V. radiata* gene | *V. radiata* protein |  |  |  |
| At5g27760 | COX-X1 |  |  | LOC106769953 | uncharacterized protein LOC106769953<sup>4</sup> |  |  |  |
| At3g43410 | COX-X5 |  |  | LOC106769343 | ARF guanine-nucleotide exchange factor GNL2 |  |  |  |
| At3g12150 | a/b hydrolase family protein |  |  | LOC106756702 | protein ABHD18 |  |  |  |
| At4g14570 | Acyl-amino-acid-releasing enzyme |  |  | LOC106772197 | acylamino-acid-releasing enzyme |  |  |  |
| At4g00960 | Putative receptor-like protein kinase |  |  | LOC106771829 | putative receptor-like protein kinase |  |  |  |

<sup>1</sup> *V. radiata* chain ID in our model.
<sup>2</sup> For subunits with multiple isoforms, only one is noted.
<sup>3</sup> Accessed on Uniprot, except for QCR10, which was accessed from EnsemblGenome with its UniParc ID.
<sup>4</sup> Identified in our mass spectrometry.

**Table 5.** Model building statistics by subunit.

| *V. radiata*<br>subunit | Uniprot ID | Chain ID | Total residues | Pre-seq. end<sup>1</sup> | Atomic residues | % Atomic<sup>2</sup> | TMH | RNA editing sites<sup>3</sup> | Ligands, lipids |
| --- | --- | --- | --- | --- | --- | --- | --- | --- | --- |
| **CIII<sub>2</sub>** |  |  |  |  |  |  |  |  |  |
| MPP-β | A0A1S3TWG4 | A | 527 | 40 | 41-527 | 100% | - | - | Zn, 2 lipids |
| MPP-α | A0A1S3VF71 | B | 506 | 49 | 50-506 | 100% | - | - | - |
| COB | E9KZM1 | C | 393 | n.a. | 2-389 | 98% | 8 | Yes | 2 x heme *b*, 9 lipids |
| CYC1 | A0A1S3UHP8 | D | 307 | 62 | 62-307 | 100% | 1 | - | heme *c*<sub>1</sub> |
| QCR1 | A0A1S3TB49 | E | 271 | 62 | 72-148 | 36% | 1 | - | See note 4 |
| QCR7 | A0A1S3U9J1 | F | 122 | n.a. | 7-122 | 94% | - | - | 1 lipid |
| QCR8 | A0A1S3U9S5 | G | 72 | n.a. | 3.0-72 | 96% | 1 | - | 4 lipids |
| QCR6 | A0A1S3VHC0 | H | 69 | n.a. | 5-69 | 93% | - | - | - |
| QCR9 | A0A1S3TQD2 | J | 72 | n.a. | 10-69 | 82% | 1 | - | 1 lipid |
| QCR10 | UPI000FCF06EC* | K | 81 | n.a. | 11-41 | 37% | 1 | - | - |
| MPP-β | A0A1S3TWG4 | M | 527 | 40 | 41-527 | 100% | - | - | Zn, 3 lipids |
| MPP-α | A0A1S3VF71 | N | 506 | 49 | 49-506 | 100% | - | - | - |
| COB | E9KZM1 | O | 393 | n.a. | 2-389 | 98% | 8 | Yes | 2 x heme *b*, 5 lipids |
| CYC1 | A0A1S3UHP8 | P | 307 | 62 | 62-307 | 100% | 1 | - | heme *c*<sub>1</sub>, 2 lipids |
| QCR1 | A0A1S3TB49 | Q | 271 | 62 | 75-148 | 35% | 1 | - | See note 4; 1 lipid |
| QCR7 | A0A1S3U9J1 | R | 122 | n.a. | 8-122 | 93% | - | - | 1 lipid |
| QCR8 | A0A1S3U9S5 | S | 72 | n.a. | 3.0-72 | 96% | 1 | - | 1 lipid |
| QCR6 | A0A1S3VHC0 | T | 69 | n.a. | 6-69 | 91% | - | - | - |
| QCR9 | A0A1S3TQD2 | V | 72 | n.a. | 11-69 | 81% | 1 | - | - |
| QCR10 | UPI000FCF06EC* | W | 81 | n.a. | 10-41 | 38% | 1 | - | - |
| Total |  |  | 4840 | n.a. | n.a. | 90% | 13 | 2 subunits | 8 ligands, 30 lipids |
| **CIV** |  |  |  |  |  |  |  |  |  |
| Cox1 | E9KZL2 | a | 527 | n.a. | 1-524 | 99% | 12 | Yes | heme *a*, heme *a*<sub>3</sub>, Cu<sub>B</sub>, Mg, 8 lipids |
| Cox2 | A0A1S3TZB2** | b | 250 | n.a. | 12-250 | 95% | 2 | - | Cu<sub>A</sub>, 4 lipids |
| Cox3 | E9KZN7 | c | 265 | n.a. | 1-265 | 100% | 7 | Yes | 6 lipids |
| Cox4 | A0A1S3TNX9*** | d | 79 | n.a. | 1-76 | 96% | 1 | - | - |
| Cox5b | A0A1S3U1E4 | e | 150 | 34 | 52-145 | 81% | - | - | Zn |
| Cox6a | A0A1S3UMA0 | f | 100 | 29 | 37-95 | 83% | 1 | - | - |
| Cox6b | A0A1S3VV60 | g | 181 | n.a. | 107-180 | 41% | - | - | - |
| Cox5c1 | A0A1S3V319 | h | 64 | n.a. | 14-62 | 77% | 1 | - | - |
| Cox7a | LOC106759053 | i | 67 | n.a. | 2-65 | 94% | 1 | - | 1 lipid |
| Cox7c | LOC106772338 | j | 96 | 29 | 47-93 | 70% | 1 | - | 1 lipid |
| Total |  |  | 1,779 | n.a. | n.a. | 88% | 26 | 2 subunits | 6 ligands, 20 lipids |

<sup>1</sup>Likely end of the mitochondrial import pre-sequence, based on homology to the corresponding *A. thaliana* subunit wherever possible or the *V. radiata* structure. n.a., not available. Note that COB and COX1-3 are mitochondrially encoded, so they do not contain import sequences.
<sup>2</sup>With respect to the number of residues of the mature protein, i.e. after cleavage of the pre-sequence. If the pre-sequence is not available, the full-length protein was used in the denominator.
<sup>3</sup>Refer to Table 5 for further details. Changes were only made when the cryoEM density was unambiguous.
<sup>4</sup>QCR1 head domain contains an iron-sulfur cluster. However, given that the head domain of QCR1 is not included in the model, the FeS cluster is not present as a ligand.
\*UniParc ID.
\*\*The *V. radiata* COX2 Uniprot entry (unreviewed, automatic entry) is mis-annotated: it contains an additional 132 N-terminal residues that are not part of the coding sequence as per the *V. radiata* genome annotation (Legume Information System, https://legumeinfo.org/gb2/gbrowse/Vr1.0). The values reported here correspond to the sequence beginning at Met133 (Uniprot numbering).
\*\*\*The *V. radiata* COX4 Uniprot entry (unreviewed, automatic entry) is mis-annotated: it contains an additional 45 N-terminal residues that are not part of the coding sequence as per the *V. radiata* genome annotation (Legume Information System, https://legumeinfo.org/gb2/gbrowse/Vr1.0). The values reported here correspond to the sequence beginning at Met46 (Uniprot numbering).

All heme groups (heme *b*_H_, *b*_L_ and heme *c*_1_) were clearly visible and modelled into each cyt *b* and cyt *c*_1_ subunits of the dimer. The distances between the hemes were consistent with that previously seen in other organisms (Figure 2B and C)^[18]^. A catalytic Zn^2+^ ion was also modelled into each MPP-*β*subunit (Figure 2B). Given that we were not able to build an atomic model for the head domain of QCR1, we did not model the iron-sulfur clusters at atomic resolution (see 3DVA analysis below for more details).

Density consistent with cardiolipin, phosphatidylethanolamine and phosphatidylcholine lipids was found, and modelled into the VrCIII_2_ map (Figure 2E). Similar to that previously seen in yeast, the CIII_2_ lipids concentrate in lipophilic cavities in COB, CYC1, QCR1, QCR8 and QCR10 close to the Q_N_ site, both on the surface of CIII_2_ and at the interface between the CIII protomers. We also observe additional lipids on the exterior of COB close to the Q_P_ site.

The vast majority of the residues that form the Q_P_ and Q_N_ sites and inhibitor-binding pockets in yeast and bovine CIII_2_ (ref. ^[15, 48, 49]^) are conserved in *V. radiata* (Figure 2-Figure Supplement 2). In particular, the COB residues that have been shown to form H-bonds with ubiquinone at the Q_N_ site of yeast and bovine CIII_2_ are conserved in *V. radiata* (His208, Ser212, D235), as are the key residues of COB’s cd1 helix (Gly149-Ile253) and the PEWY motif (Pro277-Tyr280). Nevertheless, no clear density for endogenous quinone was visible in any of the binding sites of the *V. radiata* dimer.

It is well established that the mRNAs of several subunits of plant respiratory complexes, including several CI subunits, CIII’s COB and CIV’s COX1-3, undergo cytidine-to-uridine RNA editing at several sites^[50–56]^. This mitochondrial RNA editing is widely conserved across vascular and non-vascular plants^[53, 57]^, and most generally acts to restore consensus conserved sequences^[58]^. Inspection of the cryoEM density for the VrCOB subunits showed that the *A. thaliana* COB RNA-editing sites^[59, 60]^ are conserved in *V. radiata*. Moreover, *V. radiata* COB contains additional RNA edit sites present in wheat, potato and rice^[61, 62]^ (Table 6). Given the conservation pattern and the fact that the cryoEM density at these sites was unambiguous, residues were mutated to the edited residues in our model. RNA editing was also identified in *V. radiata*’s COX1 and 3 (see CIV section), as well as CI subunits^[46]^.

**Table 6.** RNA edits identified in V. radiata CIII_2_ and CIV subunits. RNA editing sites were initially identified by an unambiguous mismatch in the cryoEM density and the expected density for the mitochondrial-DNA-encoded residue. The existence of the RNA editing site in other plants, or the implied restoration of the consensus sequence was then inspected. Amino acid changes to the atomic model (with respect to the mt-DNA sequence) were only made for amino acid positions that had unambiguous cryoEM density evidence and whose editing is conserved or would restore the conserved sequence.

| Subunit | *V. radiata* |  |  | Other species(*) |
| --- | --- | --- | --- | --- |
|  | Position | mt-DNA residue | Edited in model to: | Identical edit or consensus restoration: |
| COB | 96 | L | F | At, potato, wheat, rice<sup>[59-62]</sup> |
|  | 100 | H | Y | Potato, wheat, rice<sup>[61, 62]</sup> |
|  | 109 | H | Y | At, potato, wheat, rice<sup>[59-62]</sup> |
|  | 120 | R | W | Potato, wheat, *Chlamydomonas reinhardtii*<sup>[62]</sup> |
|  | 190 | H | Y | At, potato, wheat, rice<sup>[59-62]</sup> |
|  | 270 | P | S | Potato, wheat, rice<sup>[61, 62]</sup> |
|  | 285 | H | Y | At, potato, wheat, rice<sup>[59-62]</sup> |
|  | 303 | P | L | At, potato, wheat, rice<sup>[59-62]</sup> |
|  | 328 | H | Y | At, potato, wheat, rice<sup>[59-62]</sup> |
|  | 339 | R | C | At<sup>[59, 60]</sup> |
|  | 361 | P | S | Potato, rice<sup>[61, 62]</sup> |
|  | 375 | P | L | Potato, wheat, rice<sup>[61, 62]</sup> |
| COX1<br>(***) | 81 | S | F | (**) At, Pd, Sc, Hs, Bt |
|  | 85 | S | F | (**) At, Pd |
|  | 148 | S | L | (**) At, Pd, Sc, Hs, Bt |
|  | 151 | S | F | (**) At, Pd, Sc, Hs, Bt |
|  | 172 | S | F | (**) At |
|  | 192 | P | L | (**) At, Hs |
|  | 197 | P | L | (**) At, Sc, Hs, Bt |
|  | 223 | S | F | Variety of vascular and non-vascular plants<sup>[57]</sup> |
|  | 239 | R | W | Variety of vascular and non-vascular plants<sup>[57]</sup> |
|  | 249 | P | L | Variety of vascular and non-vascular plants<sup>[57]</sup> |
|  | 254 | S | F | Variety of vascular and non-vascular plants<sup>[57]</sup> |
|  | 396 | H | Y | (**) At, Pd, Sc |
|  | 427 | L | F | Variety of vascular and non-vascular plants<sup>[57]</sup> |
|  | 469 | R | C | (**) At |
| COX3 | 82 | P | L | At, wheat, magnolia, olive<sup>[59, 60, 87-89]</sup> |
|  | 86 | S | F | At, wheat, magnolia, olive<sup>[59, 60, 87-89]</sup> |
|  | 88 | P | L | Magnolia<sup>[89]</sup> |
|  | 102 | R | W | Magnolia, olive<sup>[88, 89]</sup> |
|  | 104 | S | F | At, wheat, magnolia, olive<sup>[59, 60, 87-89]</sup> |
|  | 105 | S | F | At, wheat, magnolia, olive<sup>[59, 60, 87-89]</sup> |
|  | 138 | P | L | At, wheat, magnolia, olive<sup>[59, 60, 87-89]</sup> |
|  | 171 | S | L | Wheat, magnolia<sup>[87, 89]</sup> |
|  | 252 | R | W | At, wheat, magnolia, olive<sup>[59, 60, 87-89]</sup> |

(\*) Non-exhaustive list.
(\*\*) RNA edit restores identity to the mt-DNA-encoded residue to the indicated model organisms (non-exhaustive): *Arabidopsis thaliana* (At), *Paracoccus denitrificans* (Pd), *Saccharomyces cerevisiae* (Sc), *Bos taurus* (Bt), *Homo sapiens* (Hs).
(\*\*\*) *A. thaliana* does not have any RNA editing sites in cox1; all the RNA editing sites identified in the other species are already mt-DNA mutations in *A. thaliana*.

#### Differences in core and non-MPP accessory subunits

Although the sequence conservation of the *V. radiata* (Vr) core CIII_2_ subunits (COB, CYC1, QCR1) with respect to *Saccharomyces cerevisiae* (Sc) and bovine (*Bos taurus*, Bt) homologues is modest (∼50%), their structural conservation is high (0.75-0.95 Å main chain RMSD, Figure 2- Figure Supplement 2-3). For QCR1, we are only able to compare the regions for which we built an atomic model, i.e. the N-terminal loop and the main helix but not the head domain. Whereas the helix is highly structurally conserved, VrQCR1 shows an extended N-terminal unstructured loop and short helix that provide more extensive contacts with MPP-*β* (see MPP section below).

Several of the accessory subunits of VrCIII_2_ show significant differences with their yeast and bovine homologues. VrMPP-*α* and -*β*, which show the most notable differences, are described in detail in the following section. Here, we discuss VrQCR7 and VrQCR8 (Figure 2-Figure Supplement 3H-J). As in mammals and yeast, VrQCR7 (ScQcr7p, BtUQCRB) contains a helix that contacts the MPP-*β* anchoring *β*-sheet (*via* VrCYC1 and VrQCR8), a helix that contacts MPP-*β* in the opposite CIII monomer and a helix that contacts COB’s surface on the matrix side (BC, DE and FG loops, and helices G and H; see Figure 2-Figure Supplement 2 for COB helix nomenclature). Nevertheless, similar to the bovine homologue, VrQCR7 is missing an N-terminal helix, in yeast, that provides several additional contacts with cyt *b*’s helix H and FG loop (Figure 2-Figure Supplement 3H). Similar to mammals and yeast, VrQCR8 (ScQcr8, BtUQCRQ) provides one strand to the multi-subunit *β*-sheet that helps anchor the MPP domain to the rest of CIII_2_ (Figure 2-Figure Supplement 3I and J). Additionally, VrQCR8’s transmembrane helix contacts VrCOB’s helices G and H. Nevertheless, like BtUQCRQ but in contrast to ScQcr8, VrQCR8 lacks a short perpendicular helix that stacks below the cyt *b*’s helix H in yeast (Figure 2- Figure Supplement 3I). Moreover, ScQcr8 also has a longer unstructured N-terminus that extends into the cavity of the MPP domain, contacting helix a and DE loop of COB in the same CIII protomer, as well as the helix a of COB of the opposite protomer. Therefore, the interaction interface between VrQCR8 and VrCOB is reduced relative to that of yeast (Figure 2-Figure Supplement 3J).

#### Differences in the MPP domain

Each CIII monomer contains an MPP-*α*/*β* heterodimer (plant MPP-*α*/*β*, yeast Cor1/2 and mammalian UQCR1/2). Given that both MPP subunits have concave surfaces that face each other, the MPP-*α*/*β* heterodimer contains a large central cavity. The VrMPP-*α*/*β*dimer shows this overall “clam shell” configuration, with a highly negative surface in the interior of the cavity (Figure 3A, Figure 3-Figure Supplement 1A-B). This negative surface has been shown to interact with the generally positively charged pre-sequences of the MPP substrates^[21]^. VrMPP-*β* contains the characteristic inverse Zn-binding HxxEH motif of pitrilysin endopeptidases, as well as all the conserved catalytic residues(Figure 3 – Figure Supplement 1)^[20]^. Our cryoEM map shows density for the Zn^2+^ ion, which is coordinated by residues His137, His141, Glu217 (Figure 3B). The fourth Zn^2+^-coordinating atom—the oxygen of the water molecule that exerts the nucleophilic attack on the carbonyl carbon of the peptide bond^[20]^—was not modelled. The glutamate that acts as a general base catalyst of this water molecule^[20]^ ^[21]^ (Glu140) is also conserved in *V. radiata*. MPP-*β* residues that were confirmed in the substrate-bound yeast soluble MPP structure^[21]^ to be critical for substrate recognition are conserved both in the sequence and structural location in VrMPP-*β* (Glu227, Asp231, Phe144, Asn167, Ala168, Tyr169; Fig 3B). Additionally, VrMPP-*α* contains the flexible glycine-rich stretch believed to be involved in substrate binding and/or product release in ScMPP-*α*^[21]^. Consistent with this conformational flexibility, our cryoEM map shows weak density for VrMPP-alpha residues 340-344.

**Figure 3.**
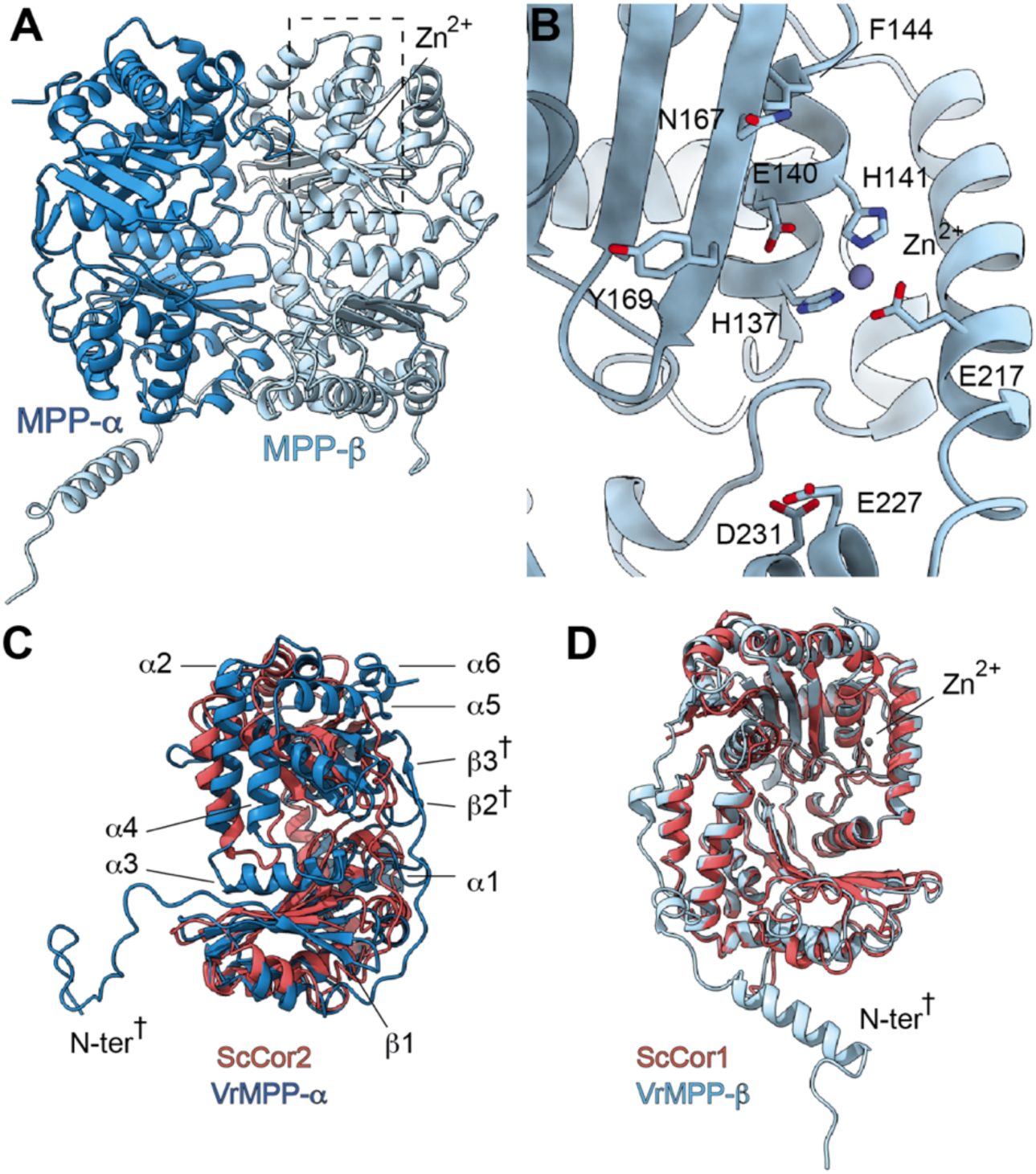
*V. radiata*’s CIII_2_ mitochondrial processing peptidase (MPP) domain has a conserved architecture and active site but contains plant-specific secondary structure elements not seen in other CIII-MPP subunits or in soluble MPP. (A) Ribbon representation of the VrMPP-*α*(blue) and VrMPP-*β* (light blue) looking into the central cavity. Dashed rectangle indicates the location of the active site, detailed in (B). (B) MPP-*β*active site [rotated 90° about vertical axis with respect to (A)]. Shown in stick representation are the Zn-coordinating residues (His137, His141, Glu217), the catalytic water-activating residue (Glu140) and conserved, putative substrate-recognition residues (Phe144, Glu227, Asp231, Asn167, Tyr169). Residue Ala168 is also conserved, but not visible in this orientation. (C-D) Superposition of *V. radiata* and *S. cerevisiae* CIII_2_ MPP domain subunits. VrMPP-*α* and -*β*’s structural elements not present in ScCor2 and ScCor1 are marked. Structural elements that are additionally not present in yeast soluble MPP, i.e. plant-specific features, are marked with a cross (^†^). (D) VrMPP-*α* (blue) and ScCor2 (dark pink). (D) VrMPP-*β* (light blue) and ScCor1 (dark pink). *β*, *β−*strand; *α*, *α*-helix; N-ter, N-terminus. See also Figure 3-Figure Supplements 1-2 for further details. *S. cerevisiae* structures from PDB: 6HU9.

VrMPP-*α*/*β*’s sequences are more similar to the yeast (and bovine) soluble MPP subunits than to the respective CIII_2_ subunits (Figure 3-Figure Supplement 1C). Consistently, the VrMPP-*α*/*β* dimer shows secondary-structure elements that are present in ScMPP-*α*/*β* and BtMPP-*α*/*β* but not in ScCor1/2. For instance, on the posterior surface of the cavity, yeast, bovine and mung bean MPP-*α* fold into six additional helices compared to ScCor2. Additionally, they contain an extra strand in the *β*-sheet underneath this helical bundle (Figure 3C, Figure 3-Figure Supplement 2).

Furthermore, some structural features of VrMPP-*α* and -*β* appear to be specific to plants, compared to the available structures (Figure 3C-D, Figure 3-Figure Supplement 1-2, Videos 4- 5). Firstly, VrMPP-*α*shows a short two-strand *β*-sheet on its posterior surface. Secondly, the extended N-terminus of VrMPP-*α* wraps over the posterior surface of VrMPP-*β*in the same CIII monomer. Thirdly, a ∼50-amino-acid N-terminal extension on VrMPP-*β* folds into an alpha helix that forms extensive contacts with MPP-*α* of the same CIII protomer as well as with VrQCR7 of the opposite protomer. A further difference in the overall configuration of the VrMPP domain comes from VrQCR1, whose longer N-terminus forms extensive plant-specific contacts with MPP-*β*’s helical bundle and *β*-sheet (Figure 3-Figure Supplement 2).

These additional contacts and secondary structure elements may serve to stabilize plants’ MPP domain. Furthermore, they strengthen the connection between the VrMPP domain and the rest of VrCIII_2_. These plant-specific features of the MPP domain could play functional roles in plant CIII_2_’s peptidase activity and, potentially, on the coupling between CIII_2_’s respiratory and non-respiratory functions (see Discussion).

#### 3D variability analysis

Given CIII_2_’s known conformational flexibility, e.g. the essential^[63–65]^ motion of QCR1’s head domain during the Q cycle, we decided to explore the conformational heterogeneity of our CIII_2_ particles using cryoSPARC’s 3D variability analysis (3DVA)^[45]^. The 3DVA algorithm uses probabilistic principal component analysis to produce distinct 3D volumes (one per principal component) that reveal the sample’s conformational heterogeneity as a continuous motion. Analyzing the individual frames of the motion of the volume, one can reconstruct discrete and continuous conformational changes of the protein sample.

We first analyzed the conformational heterogeneity of the entire CIII_2_ low-pass filtered at 6 Å. This allowed for the observation of overall changes at the level of secondary structure. Incorporating data at resolutions higher than 5 Å led to the dominance of high-resolution noise (in particular from the lipid-amphipol belt), precluding clear results. The analysis at 6 Å revealed the lack of QCR10 in some particles, as well as a change in QCR9 (principal components 0-1, Videos 6-7), indicating that these subunits only have partial occupancy in the complex. QCR10 was previously found missing in preparations of mammalian CIII_2_, likely due to de-lipidation during purification^[11, 66]^. Given that our purification is very gentle and the supercomplex interactions are maintained, it is unclear whether the changes in QCR10 and QCR9 in a sub-population of particles has biological significance or is simply due to the purification procedure. More importantly, 3DVA revealed that CIII_2_ exhibits coordinated “breathing” motions within and between the protomers of the dimer (components 0-3, Figure 4A-B, Videos 6-9). The motions, which may be parallel or anti-parallel across the dimer (compare Videos 6-8 with Video 9), extend from the top of matrix-exposed MPP domain to the bottom of the IMS-exposed regions of CYC1, QCR1 and QCR6. This finding contrasts with previous assumptions on CIII_2_’s potential for long-range conformational coupling and has implications for the potential interplay between plant CIII_2_’s respiratory and peptide-processing functions (see Discussion).

**Figure 4.**
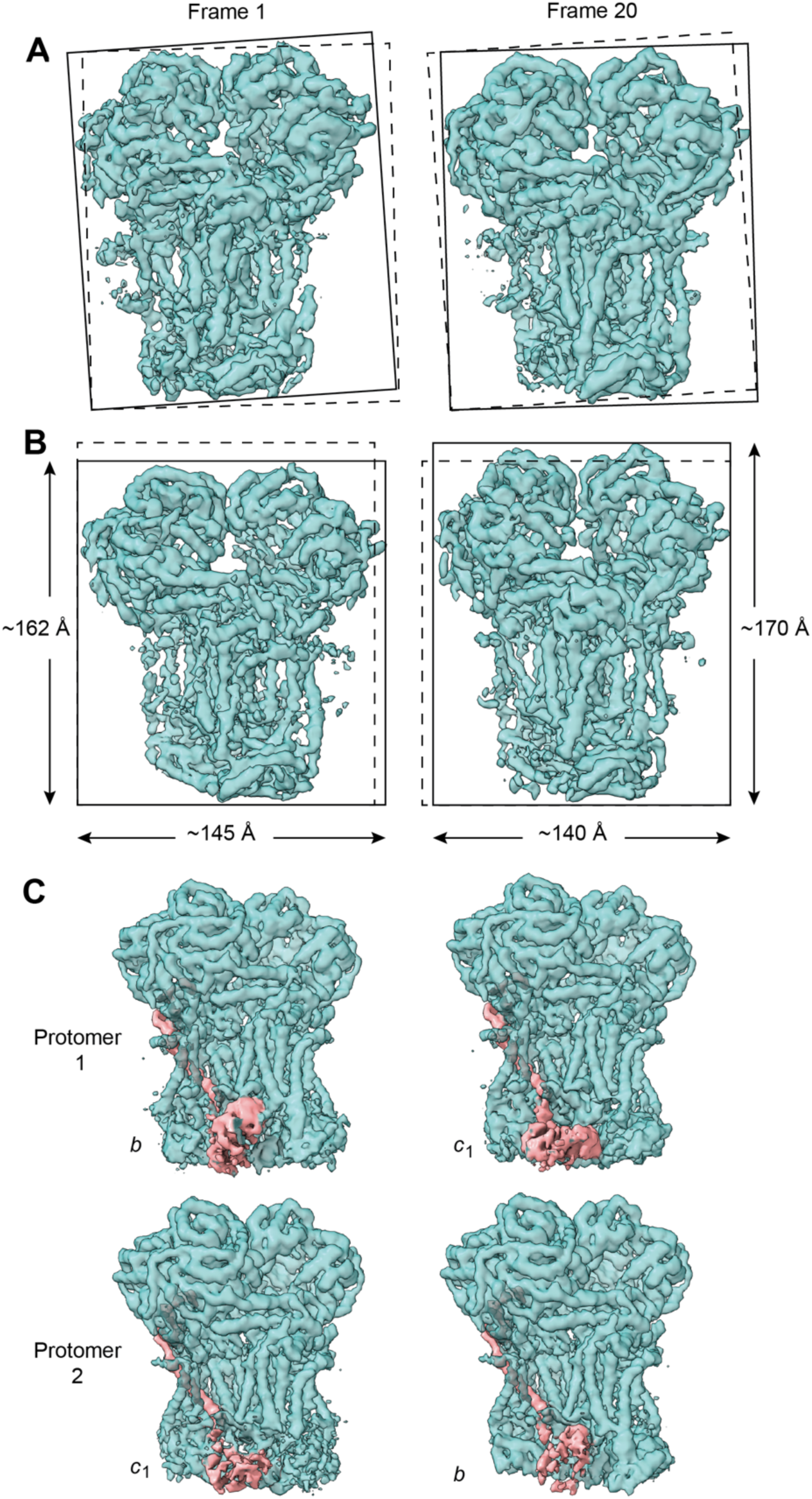
Conformational heterogeneity analysis of CIII_2_ reveals complex-wide, coordinated movements and shows the swinging motion of QCR1 in the absence of substrates or inhibitors. (A-B) CIII_2_-wide motions revealed by principal component 2 (A) and 3 (B). Frame 1 (left) and frame 21 (right) of the continuous motion of CIII_2_ (teal surface) are shown. Solid rectangles frame the outer-most edges of the complex for each frame. For comparison, dashed rectangles indicate the position of the solid rectangle of the other frame for each component. The dimensions of the rectangle sides are given in (B). (C) Frame 1 (left) and frame 20 (right) of the continuous motion of the QCR1 head domain shown for CIII_2_ protomer 1 (top) and protomer 2 (bottom). The density corresponding to QCR1 is shown in pink. The position of the QCR1 head domain is indicated by *b* (proximal) or *c*_1_ (distal). Note that, when protomer 1 is in the *b* position, protomer 2 is in the *c*_1_ position and vice versa. See Videos 6-9 for the motion of all components.

We then examined the conformational heterogeneity of the QCR1 head domain by using a mask around the IMS-exposed domains of CIII_2_ at 5 Å. The largest variability component revealed a near-continuous conformational motion in the position of the head domain of QCR1, demonstrating its swinging motion from the proximal *b* position (close to COB’s Q_P_-site) to its distal *c* position (close to CYC1) during the Q cycle^[15, 67, 68]^ (Fig. 4C). Moreover, the motions of the QCR1 head domains of the CIII dimer were anti-parallel, i.e. when one domain is in the proximal position, the other one is in the distal position and vice versa (Figure 4C, Video 10).

Given that the conformational heterogeneity precluded us from building an atomic model of the QCR1 head domain, we rigid-body fit a homology model into the extreme locations of the 3DVA volume and confirmed these corresponded to the Q-cycle proximal and distal positions (Figure 4C, Video 10). At the distal position, the FeS cluster of VrQCR1 was ∼11 Å away from the CYC1’s heme *c*_1_ (measured edge-to-edge to the ring system). This is in agreement with previous structures of the QCR1 head domain and within electron-transfer distance (14 Å)^[66, 69]^. Given that we did not model quinone in our structure, we could not measure its distance to the QCR1 FeS cluster at the proximal position. However, we measured the distance between QCR1’s FeS-coordinating residue Cys235 and COB’s Tyr285, which is within H-bonding distance to quinone^[69]^ (ScRip1-Cys180 and ScCyb-Tyr279 in yeast). This distance (∼4.5 Å) was also consistent with those previously seen^[66, 69]^, placing the *V. radiata* FeS cluster within electron-transfer distance to quinone when the head domain is at the proximal position. This confirms that the motion revealed by 3DVA is the expected swinging motion of QCR1. Moreover, given that this flexibility was observed in the absence of substrates or inhibitors, it confirms that QCR1’s head domain is intrinsically mobile.

### Complex IV

#### Overall structure and ligands

The initial resolution of CIV in SCIII_2_+IV was lower than the that of CIII_2_ (Figure 1-Figure Supplement 1). This is due to the fact that during 3D-refinement the particle poses are dominated by the larger CIII_2_ and that there is conformational flexibility at the supercomplex interface. This flexibility between CIII_2_ and CIV within SCIII_2_+IV has been previously seen in cryoEM reconstructions of CIII_2_-CIV supercomplexes in bacteria and yeast^[35–38]^. Nonetheless, focused refinements around CIV resulted in an improved reconstruction with a nominal resolution of 3.8 Å for CIV, which allowed for atomic model building (Figure 5, Figure 1-Figure Supplement 2, Video 2).

**Figure 5.**
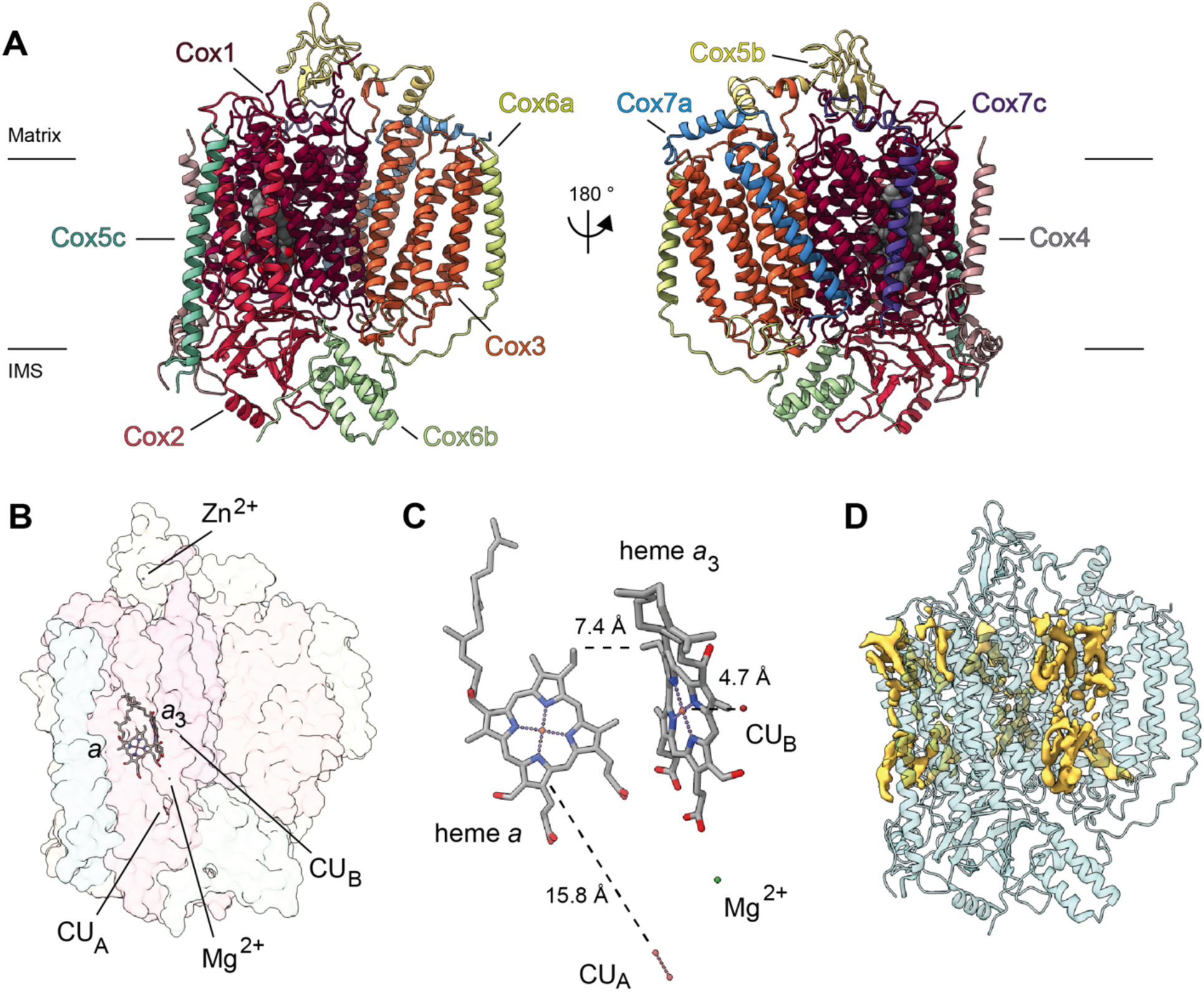
Overview of *V. radiata* CIV and its co-factors. (A) CIV in cartoon representation colored by subunit with co-factors in sphere representation colored by atom. The position of the inner mitochondrial membrane is indicated with black lines, and matrix and inter-membrane space (IMS) are labelled. (B) Position of the observed CIV co-factors. CIV shown in transparent surface representation, cofactors in stick representation. (C) Edge-to-edge distances between the heme groups and the copper co-factors are shown. The co-factors are rotated 20 degrees relative to (B) for clarity. (E) The positions of the modelled lipids (sticks) are shown on CIV (transparent surface). Cardiolipin in orange, phosphoethlyethanolamine in yellow. *a*, heme a; *a*_3_, heme *a_3_*.

At this resolution, all prosthetic groups (heme *a*, heme *a*_3_, dinuclear copper A and copper B at the binuclear center), as well as a Mg^2+^ ion and a Zn^2+^ ion were visible and modelled into the structure (Figure 5B-C). The residues that coordinate the prosthetic groups, including the covalent bond between Cε-Tyr247 and the Nε-His243 that coordinates copper B in VrCOX1, are clearly visible and hence conserved in *V. radiata*. The distances between the prosthetic groups are consistent with those seen in other organisms (Figure 5C)^[27]^. Additionally, density consistent with cardiolipin, phosphatidylethanolamines and phosphatidylcholines was seen and modelled into the CIV map (Figure 5D). These lipids and acyl chains are located in hydrophobic cavities of VrCOX1 and VrCOX3, in similar locations to those seen in yeast and bovine CIV. Moreover, conserved RNA editing sites were unambiguously identified in VrCOX1 and VrCOX3 and thus changed in the model (Table 6).

#### Plant CIV composition

The subunit composition of plant CIV has remained unclear, with different mass spectrometry studies suggesting 5-11 subunits, including some putative plant-specific subunits^[70–72]^. Our structure shows that VrCIV is composed of 10 subunits (3 core and 7 accessory). In contrast, mammalian CIV contains 11 accessory subunits and yeast CIV contains 9. Thus, although VrCIV’s overall architecture is similar to other organisms, given the lack of certain homologues (ScCox26, ScCox6/BtCOX5a, BtCOX7b, BtCOX8, BtNDUFA4), there are significant differences (Figure 6, Figure 6-Figure Supplement 1).

**Figure 6.**
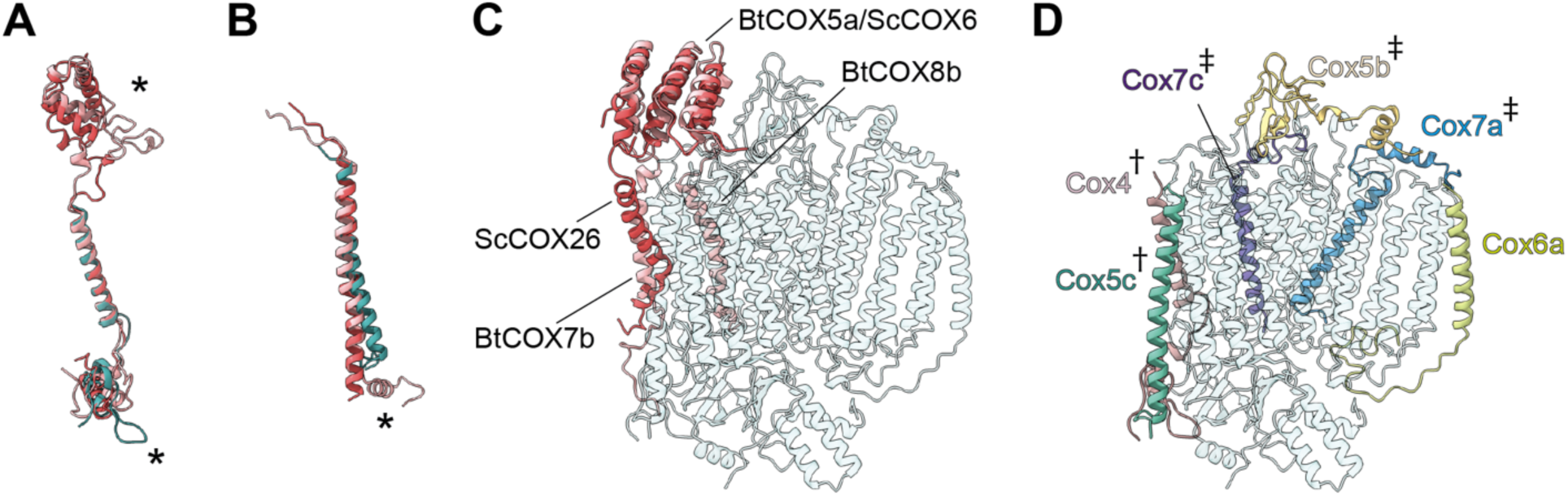
Subunit differences in CIV. (A-C) Superposition of subunits of *V. radiata* (Vr, teal), *S. cerevisiae* (Sc, dark pink; PDB: 6HU9) and *B. taurus* (Bt, light pink; PDB: 5B1A). Subunits were aligned to the corresponding *V. radiata* subunit. Asterisks mark the differences discussed in the text. (A) Superposition of VrCOX4, ScCox5a, BtCOX4. (B) Superposition of VrCOX5c, ScCox9, BtCOX6c. (C) Superposition of the yeast and bovine CIV subunits that do not have homologues in V. radiata, onto the V. radiata CIV model (transparent teal). Alignment by COX1 subunits. (D) Location of the *V. radiata* accessory subunits that show notable differences with their yeast/bovine homologues. Subunits that form the supercomplex interface in *V. radiata* are marked with (†). Subunits whose homologues form the supercomplex interface in the *B. taurus* respirasome are marked with (‡).

The presence of the *V. radiata* CIV subunits in the structure was confirmed by mass spectrometric analysis of our cryoEM sample (Table 1). Our structure confirms the identity of plant COX-X2 as the homologue of mammalian COX4 (ScCox5), COX-X3 of mammalian COX7c (ScCox8) and COX-X4 of mammalian COX7a (ScCox7)^[71]^. The putative plant-specific subunits COX-X5, GLN2, ABHD18, AARE and PRPK^[70, 71]^ were not observed in our mass spec sample or our structure (Table 1). Although two peptides of the putative subunit COX-X1 was identified in our mass spec sample, the lack of unassigned density in our cryoEM reconstruction large enough to constitute an additional subunit demonstrates that this protein is not present in SCIII_2_+IV of etiolated *V. radiata* tissues. The possibility remains that free CIV and/or that in supercomplexes of non-etiolated tissues may have a different subunit composition.

#### Differences in core and accessory subunits

Among the VrCIV subunits that have mammalian and yeast homologues, structural conservation is generally high, particularly for the core subunits (Figure 5-Figure Supplement 1). The conserved covalent bond between Nε-His243 and Cε-Tyr247 on the HPEVY ring of VrCOX1 is clearly seen in our density (Fig 5-Figure Supplement 1J). Moreover, VrCOX2 contains the highly conserved residues Tyr255, Met-362 (BtCOX2-Tyr105, Met207) believed to be part of the electron transport path from cyt *c* to CU_A_ of CIV^[73]^.

A significant difference is seen in VrCOX4, which is missing a ∼100-amino-acid N-terminal helical domain compared to its homologues (ScCox5, BtCOX4; Figure 6A and C). In yeast and mammals, this N-terminal helical domain mainly interacts with ScCox6/BtCOX5a, which is one of the accessory subunits that is missing in plants (Table 4). Moreover, in yeast SC III_2_+IV_2_, this helical domain of ScCox6 interacts with MPP-β of CIII_2_ to provide the main CIII:CIV contacts for supercomplex formation. Hence, this interface is absent in the plant SC III_2_+IV (see SC section below).

Further differences are found in VrCOX4’s C-terminus, which shows an extended loop that reaches towards the IMS side of VrCOX2 and then folds back towards the solvent-accessible face of CIV. Another difference in the vicinity of VrCOX4 is seen in VrCOX5c, which lacks the C-terminal helix that in BtCOX6c makes additional contacts to BtCOX2 and BtCOX4 (Figure 6A, Figure 6-Figure Supplement 1). These differences in VrCOX4 and VrCOX5c—and the lack of a homologue for ScCox5/BtCOX4—are notable, as these subunits form the majority of interactions between CIV and CIII_2_ in the plant SC III_2_+IV (see SC section below). Differences in VrCOX5b (shorter N-terminus), VrCOX7a (longer N-terminus with additional contacts to VrCOX3) and VrCOX7c (different path for the unstructured N-terminus) compared to their mammalian counterparts (Figure 6-Figure Supplement 1) are also likely related to the fact that these subunits provide SC-interface contacts in mammals not observed in plants. However, whether these differences are a cause or an effect of the divergent supercomplex binding interfaces remains to be determined.

Additional differences are seen in VrCOX6a (ScCox13, BtCOX6a). In this case, the plant subunit is more similar to the bovine homologue. Its shorter helix interacts with VrCOX1 and VrCOX3 on the IMS side but does not provide matrix-side interactions as the yeast homologue does (Figure 6-Figure Supplement 1).

#### CIV’s proton transfer pathways

Translocation of protons through CIV in different organisms occurs via the D, K and H “channels” (proton transfer pathways) of the COX1 subunit^[27–30]^. The D and K channels are essential for coupled proton transfer in CIV of bacteria, yeast and mammals^[27–31]^. However, whereas mutations of key H channel residues abolish proton pumping in bovine CIV^[74, 75]^, analogous mutations have no effect on growth, respiratory rate or proton-to-electron ratios of yeast CIV (ref. ^[31]^). Thus, it is currently thought that the H channel only plays a role in proton transfer in mammalian CIV (although it may still act as a dielectric channel in yeast^[28, 31]^). The contribution of the D, K and H channels to CIV proton transfer in plants is unknown.

Sequence and structural analyses show that all protonatable residues of the D and K channels of yeast and mammals are conserved in VrCOX1 (Figure7, Figure 7-Figure Supplement 1). In contrast, several of the residues of the mammalian H channel are not conserved in mung bean (Figure 7-Figure Supplement 1). In bovine’s H channel, the amide bond between Tyr440 and Ser441 and a H-bond network towards Asp51 are essential features for proton translocation^[29, 74, 76]^. None of these key bovine residues are conserved in *V. radiata* (or *S. cerevisiae*) (Figure 7, Figure 7-Figure Supplement 1). Moreover, in the bovine H channel mutation of Ser441 to Pro441 abolishes proton pumping^[74]^. The corresponding amino acid for BtCOX1-Ser441 in both *V. radiata* and yeast is proline (VrCOX1-Pro443; ScCox1-Pro441). Additional amino acid differences between *V. radiata* and *B. taurus* are seen at the entrance and exit of the H channel (Figure 7). The above suggests that, similar to yeast, the H channel is not a proton transfer pathway in plant CIV and that proton transfer in plant CIV occurs *via* the D and K channels. Additional, experimental evidence is needed to confirm this hypothesis.

**Figure 7.**
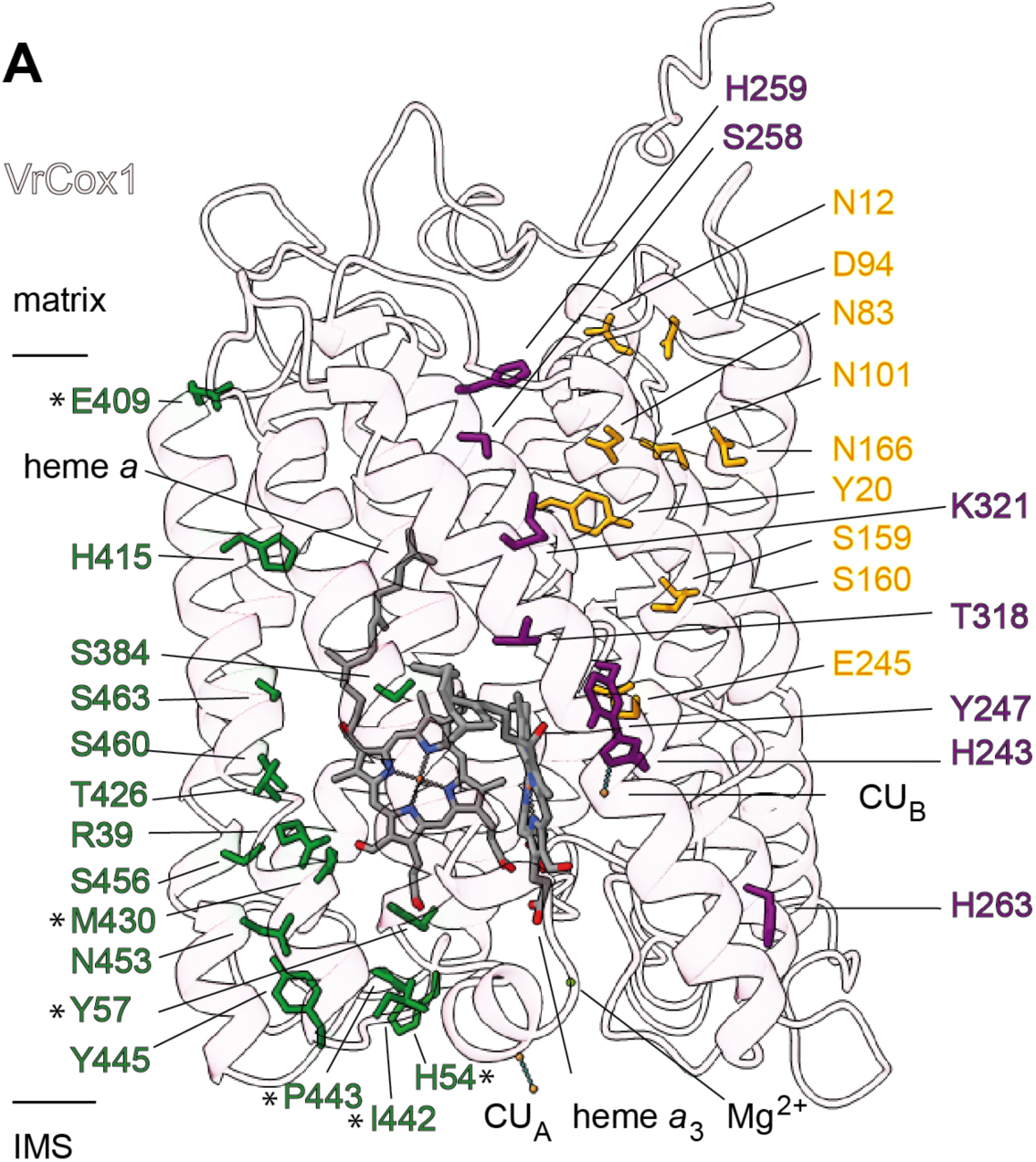
Proton transfer pathways of *V. radiata* CIV. VrCOX1 (transparent ribbon), co-factors (stick) and key residues (colored stick) are shown for the D channel (yellow), K channel (purple) and H channel (green). Proton-channel residues that are mutated in *V. radiata* with respect to *B. taurus* are marked with an asterisk (*). Approximate position of matrix and IMS ends of the transmembrane region are shown.

### Supercomplex III_2_-IV (SC III_2_+IV)

By docking the individually refined CIII_2_ and CIV models into the SC III_2_+IV composite map (Figure 1, Figure 1-Figure Supplement 2, Video 3), we were able to define the binding interface of the plant supercomplex. Direct contacts between the complexes occur in one site in the matrix side and one site in the IMS (Figure 8). Site 1 (matrix side), shows a single hydrophobic contact between one residue of VrQCR8 (Pro31) and one residue of VrCOX2 (Trp59) (Fig. 8B). Our cryoEM reconstruction also contains a short stretch of weak, unassigned density near the first modelled residue of VrCOX5c (BtCOX6c, ScCox9), which could maximally represent an additional four amino acids. If so, the N-terminus of VrCOX5c could potentially provide additional contacts with CIII_2_. However, this density may also correspond to a bridging lipid bound between the two complexes.

**Figure 8.**
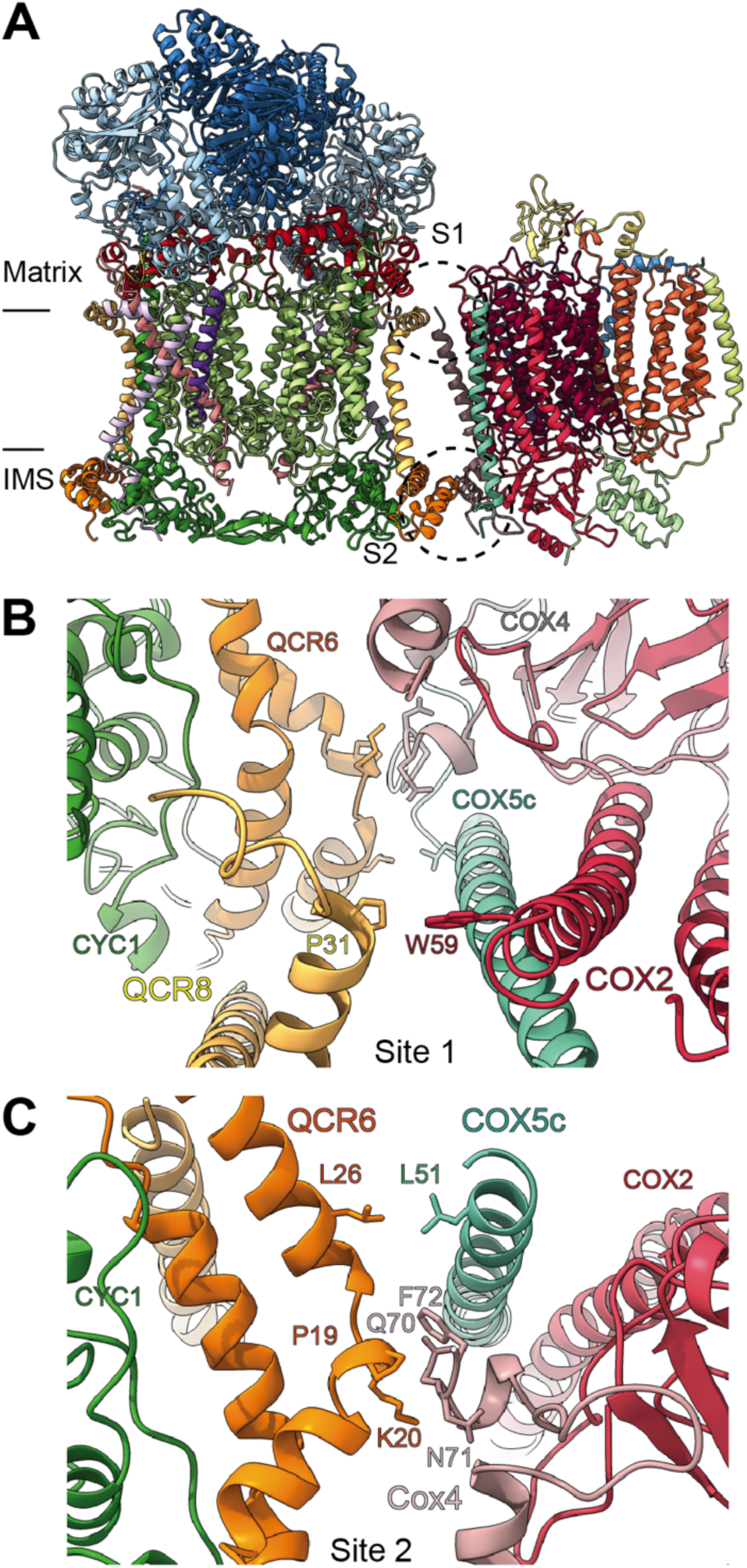
SCIII_2_+IV interface in *V. radiata*. (A) General orientation of SCIII_2_+IV in ribbon representation viewed from the membrane. Approximate position of the inner mitochondrial membrane is shown. Sites 1 (S1) and 2 (site 2) of the supercomplex interface are marked in dashed circles. (B) Detailed view of the protein-protein interaction in site 1 (Pro31 of VrQCR8 and Trp59 of VrCOX2) with the interacting atoms shown in stick representation. Note that interacting residues of site 2 appear in stick in the background. (C) Detailed view of the protein-protein interaction in site 2 (Pro19-Lys20 of QCR6, Gln70-Phe72 of COX4, Leu26 of QCR6, Leu51 of COX5c) with the interacting atoms shown in stick representation.

The limited matrix-side contacts in *V. radiata* is in stark contrast to the supercomplex interface in *S. cerevisiae*, where binding is dominated by interactions between ScCor1 (VrMPP-β) and ScCOX5a (VrCox4) on the matrix side. Instead, *V. radiata*’s site 2 (IMS side) provides the bulk of the protein-protein interactions of the supercomplex, with a hydrophobic interaction between VrQCR6 and VrCOX5c (Leu26 and Leu51 respectively) as well as an interface between VrQCR6 (Pro19 and Lys20) and VrCOX4 (Arg114-Phe117) (Fig. 8C). Despite the potential for lipid bridges at the matrix leaflet of the IMM, there are no direct contacts in the membrane and no protein contact at the IMS leaflet of the IMM.

Unsurprisingly given the large differences in contacts, there is a significant difference in the angle between CIII_2_ and CIV in *V. radiata versus* yeast, resulting in a more “open” orientation in *V. radiata* (Figure 8-Figure Supplement 1). This orientation leads to a difference of 18° in the angle between the CIII_2_ and CIV as measured by the relative positions of the *b*_h_-hemes in CIII_2_ and the *a*-hemes in CIV. The difference in orientation also results in a larger estimated distance between the CIII_2_- and CIV-bound cyt *c* in *V. radiata* (∼70 Å) than in yeast (∼61 Å) (Figure 8- Figure Supplement 1E).

## Discussion

Here, we present the first high-resolution structures and atomic models of CIII_2_, CIV and supercomplex SC III_2_+IV from the plant kingdom (Figure 1, Videos 1-3). These *V. radiata* structures reveal atomic details of the catalytic sites and co-factor binding of plant CIII (Figure 2) and CIV (Figure 5). Moreover, they show plant-specific structural features for several subunits, most notably the MPP subunits of CIII_2_ (Figure 3). Conformational heterogeneity analysis of CIII_2_ allowed us to observe the swinging motion of QCR1’s head domain in the absence of substrates or inhibitors, and revealed coordinated, complex-wide motions for CIII_2_ (Figure 4, Videos 6-10). The CIV structure defines the subunit composition of the plant complex and suggests that the proton translocation in plants occur via the D- and K-, rather than the H-channel (Figures 5, 6 and 7). The structures also reveal the plant-specific arrangement of the CIII_2_-CIV interface in the supercomplex, which occurs mostly on the IMS side of the membrane and at a different angle from that seen in other organisms (Figure 8 and Figure 8-Figure Supplement 1).

### Conformational heterogeneity of plant CIII_2_

The 3DVA allowed us, for the first time, to observe the mobility of the QCR1 (Rieske subunit) head domain in a single preparation, without the need for CIII_2_ inhibitors to induce the proximal or distal position (Figure 4, Video 10). As such, it provides direct confirmation for previous crystallographic, mutational, kinetic and molecular dynamics studies^[15, 67, 68, 77]^, showing that the flexible swinging motion is an inherent property of the QCR1 in the absence of substrates. Moreover, the conformational heterogeneity analysis showed that the movement of the QCR1 head domains is coordinated in an antiparallel fashion between the CIII_2_ protomers (Figure 4C, Video 10).

Unfortunately, however, this conformational flexibility precluded us from building an atomic model for the head domain. Thus, we were not able to evaluate the H-bonding pattern of the QCR1 head domain with COB, or the implications of such pattern to the Q-cycle electron bifurcation mechanism^[15, 51, 52]^. Similarly, the resolution of the 3DVA was not sufficient to evaluate changes in the positions of COB’s cd1 and ef helices. These helices are critical components of the QCR1’s COB binding “crater”^[15]^, and their location changes in response to binding of different CIII_2_ inhibitors^[78]^, with important implications for the Q-cycle mechanism. It is important to note that 3DVA only reveals conformational changes, with no information on kinetics or occupancy rates. Nonetheless, we demonstrate here that cryoEM conformational heterogeneity tools such as 3DVA^[45]^ and others^[79]^ are a valuable complementary approach to study the conformational changes of CIII_2_’s Q-cycle in its native state, as well as in the presence of inhibitors and substrates.

Moreover, the 3DVA revealed that CIII_2_ can undergo different types of complex-wide motions that are coordinated across sides of the membranes and between protomers (Videos 6-9). Changes at the “top” of the complex on one side of the membrane co-vary (i.e. are coupled) with movement at the “bottom” of the complex on the other side of the membrane. This long-range conformational coupling across the entire CIII_2_ could be the basis for symmetry-breaking and coordination of the QCR1 head domain motion between the CIII_2_ protomers.

The long-range conformational coupling is particularly relevant in the context of plant CIII_2_’s dual roles in signal peptide processing and respiration. The potential interdependence between these two functions has been investigated using CIII_2_ inhibitors^[22, 26]^. In the presence of the CIII_2_ respiratory inhibitors antimycin A (Q_N_-site inhibitor) and myxothiazol (Q_P_-site inhibitor) at concentrations that inhibit spinach CIII_2_’s respiratory activity ∼90%, the complex’s peptidase activity was inhibited 30%-40%^[22, 26]^. At the time, in the absence of any structural information for any CIII_2_, these results were interpreted as suggesting that inhibition of the peptidase activity of plant CIII_2_ is not a result of the inhibition of the respiratory activity of the complex. When crystal structures of metazoan CIII_2_ in complex with these inhibitors became available and revealed the locations of the inhibitor binding sites^[14, 16, 19]^, the large distances between these sites and the MPP domain were interpreted to reinforce the notion that the dual roles of plant CIII_2_ were independent, as long-range coupled conformational changes were deemed unlikely^[23]^. In contrast, our 3DVA results showed that long-range coupled motions are intrinsic to *V. radiata* CIII_2_. Moreover, our atomic model of plant CIII_2_ revealed additional contacts and secondary structure elements not previously seen in other organisms that enhance the interaction between the MPP domain and the rest of the complex. For example, the extended N-termini of MPP-*α* and -*β*bridge across the dimer and provide plant-specific contacts with CIII_2_’s membrane subunits (Figure 3, Figure 3-Figure Supplements 1-2). Moreover, QCR1’s longer N-terminus in plants also provides plant-specific contacts with the MPP domain (Figure 2-Figure Supplements 1, 3; Figure 3-Figure Supplement 1C). Given its span across the membrane and its domain-swapping across protomers, QCR1 may have roles as a “conformational coupler” beyond its essential function in the Q-cycle.

Together, the 3DVA results challenge long-standing assumptions on plant CIII_2_’s suitability for conformational coupling and call for a re-evaluation of the relationship between the respiratory and the processing activities of the plant complex.

### Plant supercomplex III_2_+IV interface

The orientation and binding interfaces of SC III_2_+IV vary significantly among organisms^[35–38, 80]^. For instance, the CIII surface used by yeast to bind to CIV is instead used by mammals to bind to CI^[35]^. Given these disparities, it is not surprising that the differences seen in the VrCIV subunits are concentrated in the subunits that form the supercomplex interfaces in the different organisms (Figure 6). Moreover, while some of the supercomplex interactions in *V. radiata* are reminiscent of the supercomplex interface in yeast, there are significant differences in the protein:protein sites, interacting subunits and angle of orientation within the SC (Figure 8 and Figure 8-Figure Supplement 1). In yeast, the main interface is on the matrix side, with the N-terminal helical domain of ScCox5a (VrCOX4) interacting with ScCor1 (homologue of VrMPP-β). In *V. radiata*, this interface is lacking, as plant COX4 does not possess the ∼100-amino-acid helical N-terminal domain present in yeast and mammals (Figure 6 and Figure 6-Figure Supplement 1). In contrast, the main supercomplex interface in mung bean is in the IMS, driven by contacts between VrQCR6 and VrCOX4. In the yeast supercomplex, the homologues of VrQCR6 and VrCOX4 (ScQCR6 and ScCOX5a/b) also interact in the IMS side, but in a much more limited fashion.

In light of VrCIII_2_’s conformational heterogeneity and the potential interdependence between respiratory and peptidase functions of plant CIII_2_, an intriguing possibility is that matrix-side interactions between CIII_2_ and CIV are minimized in the plant supercomplex to prevent steric constraints on the MPP domain, which is catalytically active and likely requires flexibility for its peptidase activity. Thus, the plant-specific supercomplex interface may have evolved to accommodate the particularities of plant CIII_2_’s dual respiratory and processing functions.

Nevertheless, whereas the details differ, the overall location of the CIII_2_:CIV interfaces in *V. radiata* and yeast is similar. A related observation has been made for the supercomplexes between CIII_2_ and CI (SCI+III_2_) of plants, yeast and mammals as seen by sub-tomogram averaging^[39]^. In this case, the interfaces between CI and CIII_2_ in the different organisms were also similar; however, there was a ∼10° difference in the angle between CI and CIII_2_. Additional functional/structural studies of supercomplexes from organisms of diverse phylogenetic origins could determine whether the location of the supercomplex interface have been achieved by convergent or divergent evolution. In turn, this would shed light on the evolution and potential functional significance of the interface sites.

What can already be concluded is that—as seen in yeast^[35, 36]^—the benefit of the SC III_2_+IV arrangement in plants does *not* involve direct electron transfer from CIII_2_ to CIV by simultaneously bound cyt *c* on each complex, as the calculated distance between the bound cyt *c* is too large (∼70 Å, Figure 8-Figure Supplement 1). Recent quantitative-proteomics estimations of the stoichiometry of plant respiratory chain components indicate that the average plant mitochondrion contains ∼6,500 copies of CIII monomers (i.e. ∼3,250 CIII_2_), ∼2,000 copies of CIV and ∼2250 copies of cytochrome *c*^[81]^. This implies a maximum of ∼2,000 copies of SC III_2_+IV and, thus, a roughly 1:1 ratio between cyt *c* and SC III_2_+IV. (The ratio has been estimated to be 2-3 in *S. cerevisiae*^[82]^). Based on recent theoretical analyses of electron flow between CIII_2_ and CIV^[82]^, at this low 1:1 ratio, electron flow would be limited by the time constant of cyt *c* equilibrating with the bulk IMS phase. Under these conditions, the formation of SC III_2_+IV in the plant mitochondrion would provide a kinetic advantage to electron flow between CIII_2_ and CIV by reducing the distance between them relative to CIII_2_ and CIV freely diffusing in the plane of the membrane. It is important to note that this possible kinetic advantage does not imply substrate trapping or channeling between the complexes, and is thus consistent with a single cyt *c* pool^[82]^.

This work provides the first high-resolution structure of SC III_2_+IV in plants, revealing plant-specific features of the complexes and supercomplex. This will allow for the development of more selective inhibitors for plant CIII and CIV, frequently used as agricultural herbicides and pesticides^[83]^. The structures also allow for the generation of new mechanistic hypotheses—for example, related to proton translocation in CIV—and a re-evaluation of long-standing assumptions in the field—for instance, related to CIII’s capacity for long-range coordinated motion and the relationship between its respiratory and processing functions. Together with biochemical, cellular and genetic studies, further comparative analyses of these atomic structures with the growing number of respiratory complexes and supercomplexes across the tree of life will allow for the derivation of the fundamental principles of the respiratory electron transport chain.

## Video legends

Video 1. CryoEM density map and model for *V. radiata* CIII_2_.

Video 2. CryoEM density map and model for *V. radiata* CIV.

Video 3. CryoEM density map and model for *V. radiata* SC III_2_+IV.

Video 4. Superposition of *V. radiata* MPP-*β* with S. cerevisiae Cor1 (6HU9) and soluble MPP-*β*(1HR6).

Video 5. Superposition of *V. radiata* MPP-*α* with S. cerevisiae Cor2 (6HU9) and soluble MPP-*α*(1HR6).

Video 6. 3D variability analysis of *V. radiata* CIII_2_, component 0. The 3DVA volumes are shown as a continuous movie. CIII_2_ in teal, QCR10 in purple.

Video 7. 3D variability analysis of *V. radiata* CIII_2_, component 1. The 3DVA volumes are shown as a continuous movie. CIII_2_ in teal, QCR9 in lilac, QCR10 in purple.

Video 8. 3D variability analysis of *V. radiata* CIII_2_, component 2. The 3DVA volumes are shown as a continuous movie. CIII_2_ in teal.

Video 9. 3D variability analysis of *V. radiata* CIII_2_, component 3. The 3DVA volumes are shown as a continuous movie. CIII_2_ in teal.

Video 10. Swinging motion of the *V. radiata* QCR1 head domains. The 3DVA volumes are shown as a continuous movie. A *V. radiata* QCR1 head-domain homology model was rigid-body fit into the 3DVA volume.

---

**Figure 1- Figure Supplement 1.**
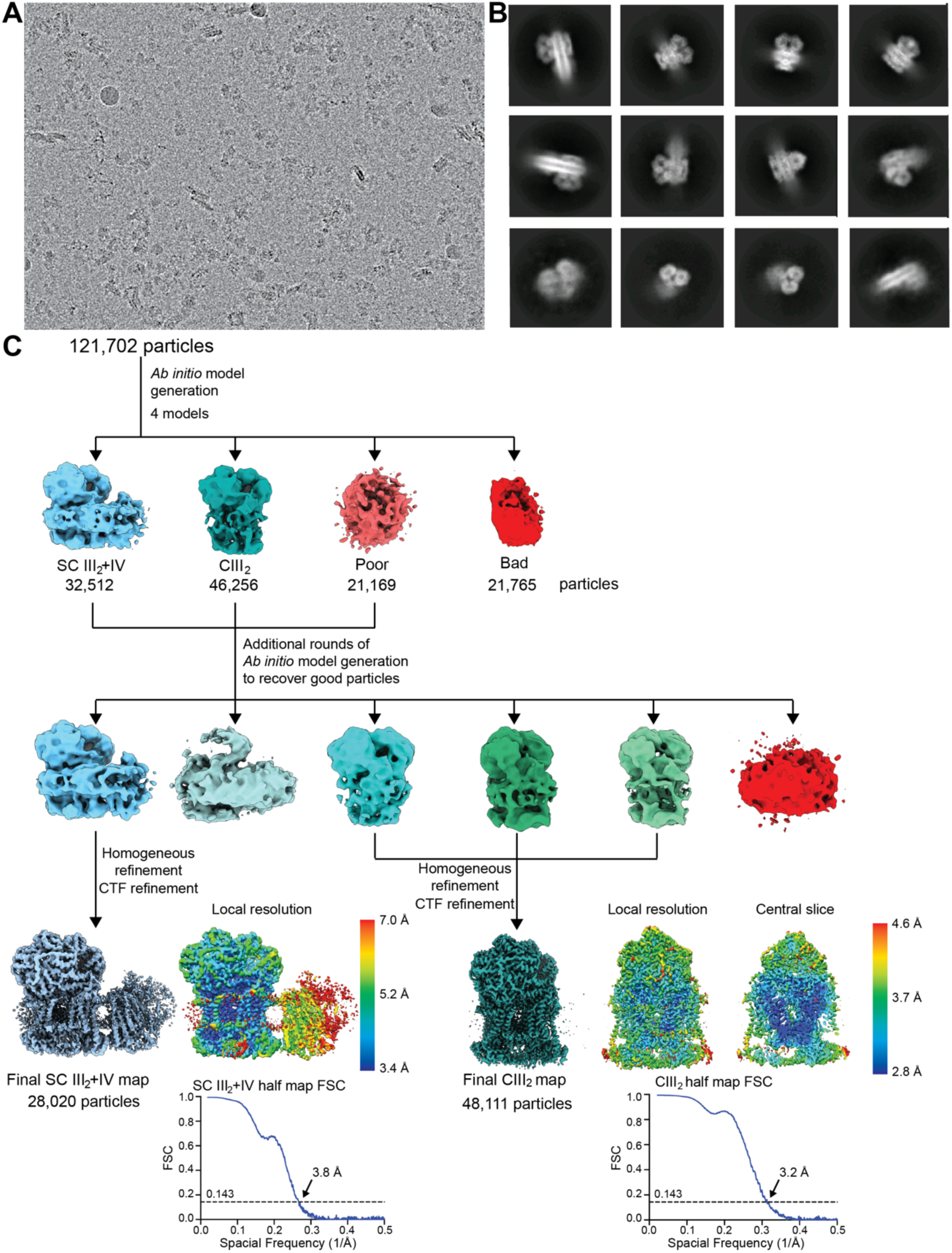
Initial processing and reconstructions using cryoSPARC. (A) A representative micrograph of the 8,541 used for further processing (9,816 collected). (B) Representative 2D class averages from reference-free classification. (C) Classification and refinement procedures used. The local resolution map and the half map gold-standard FSC curves are shown next to their respective final reconstructions.

**Figure 1-Figure Supplement 2.**
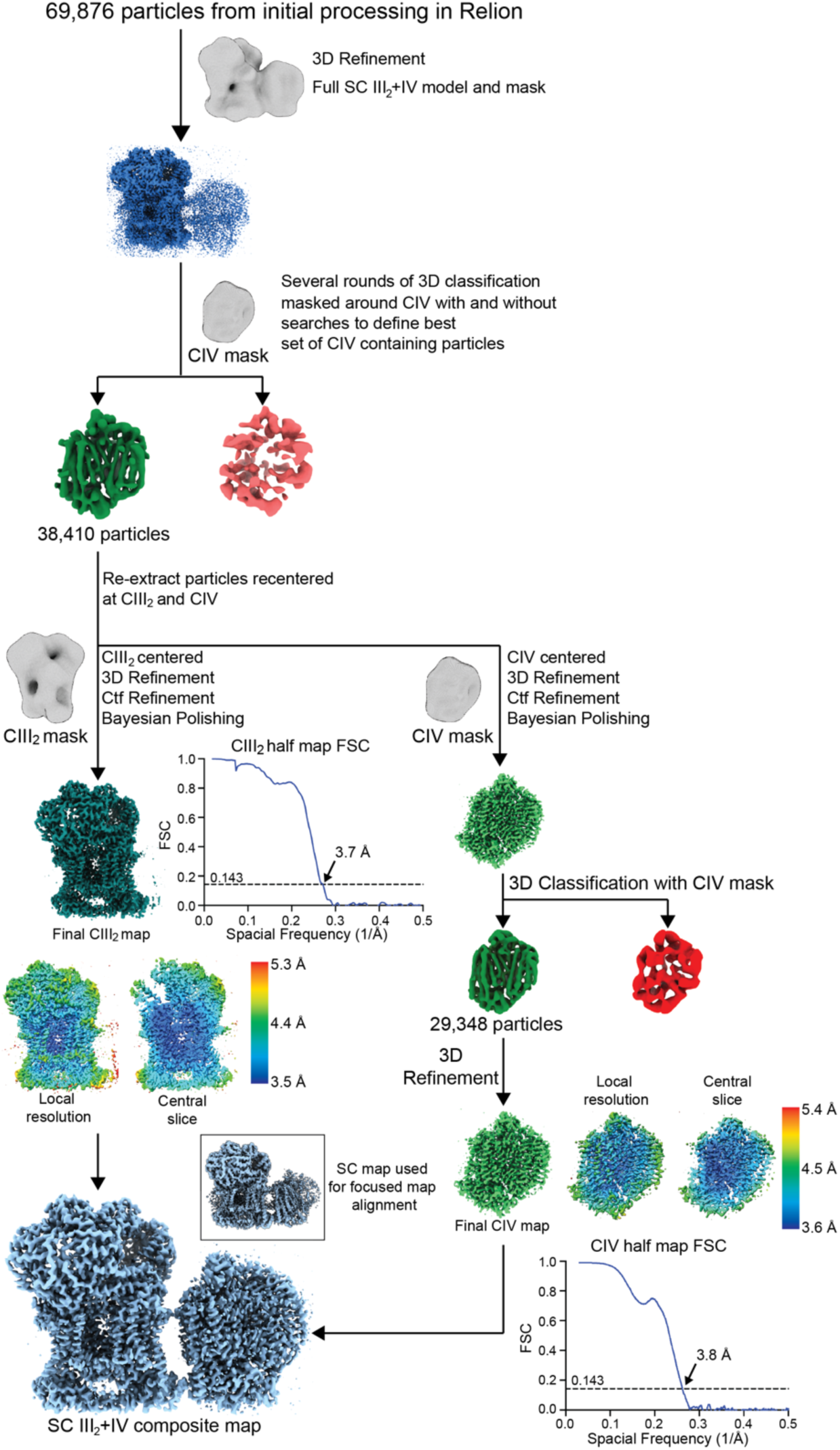
Supercomplex focused classification and 3D refinement using Relion. SC III2+IV particles selected from 2D classification were first aligned using a SC III2+IV model. These aligned particles were sorted by several rounds of 3D classification using a CIV-only mask (see Methods). Results from the 3D classifications were pooled and duplicate particles were. removed. Particles from this set were re-extracted centered at CIII_2_ and CIV, generating two sets of re-centered particles. These were 3D-refined, CTF-refined and polished independently using a CIII_2_ and CIV mask/model respectively. For the CIV-centered particles, an additional round of 3D classification was performed on the shiny particles. The local resolution map, a slice through the local resolution map and the half map gold-standard FSC curves are shown next to their respective final reconstructions. The locally refined maps were combined in Phenix to generate a composite map based on the best SC III_2_+IV reconstruction (bottom left inset; see also Figure 1-Figure Supplement 1).

**Figure 1-Figure Supplement 3.**
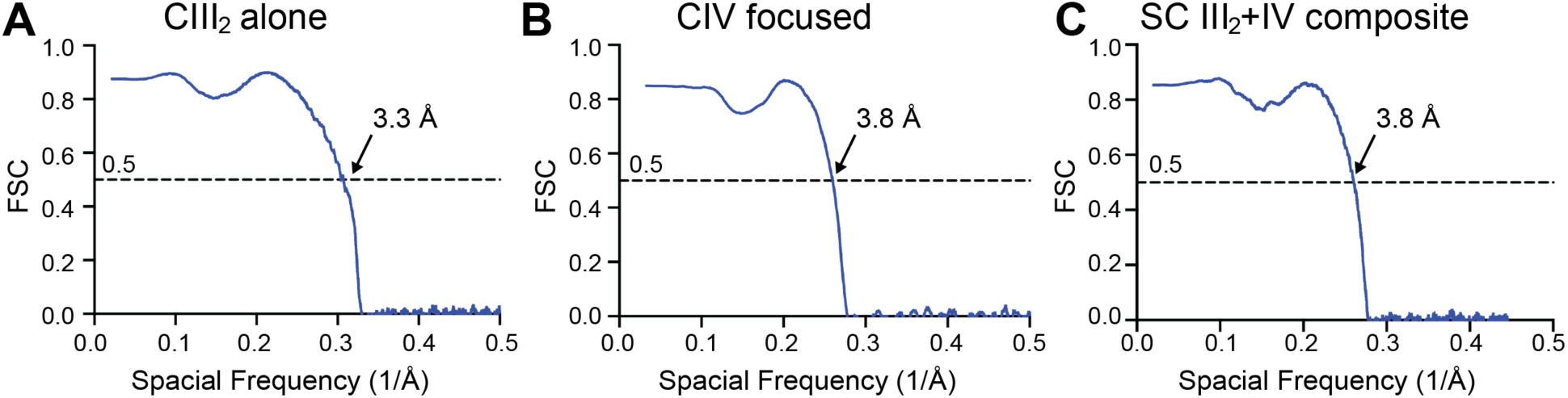
Map-Model FSCs are shown for (A) CIII_2_ alone (see also Figure 1-Figure Supplement 1), (B) CIV-focused map from the supercomplex particles (see also Figure 1-Figure Supplement 2) and (C) the SC III_2_+IV composite map (see also Figure 1-Figure Supplement 2). See also Table 3.

**Figure 2-Figure Supplement 1.**
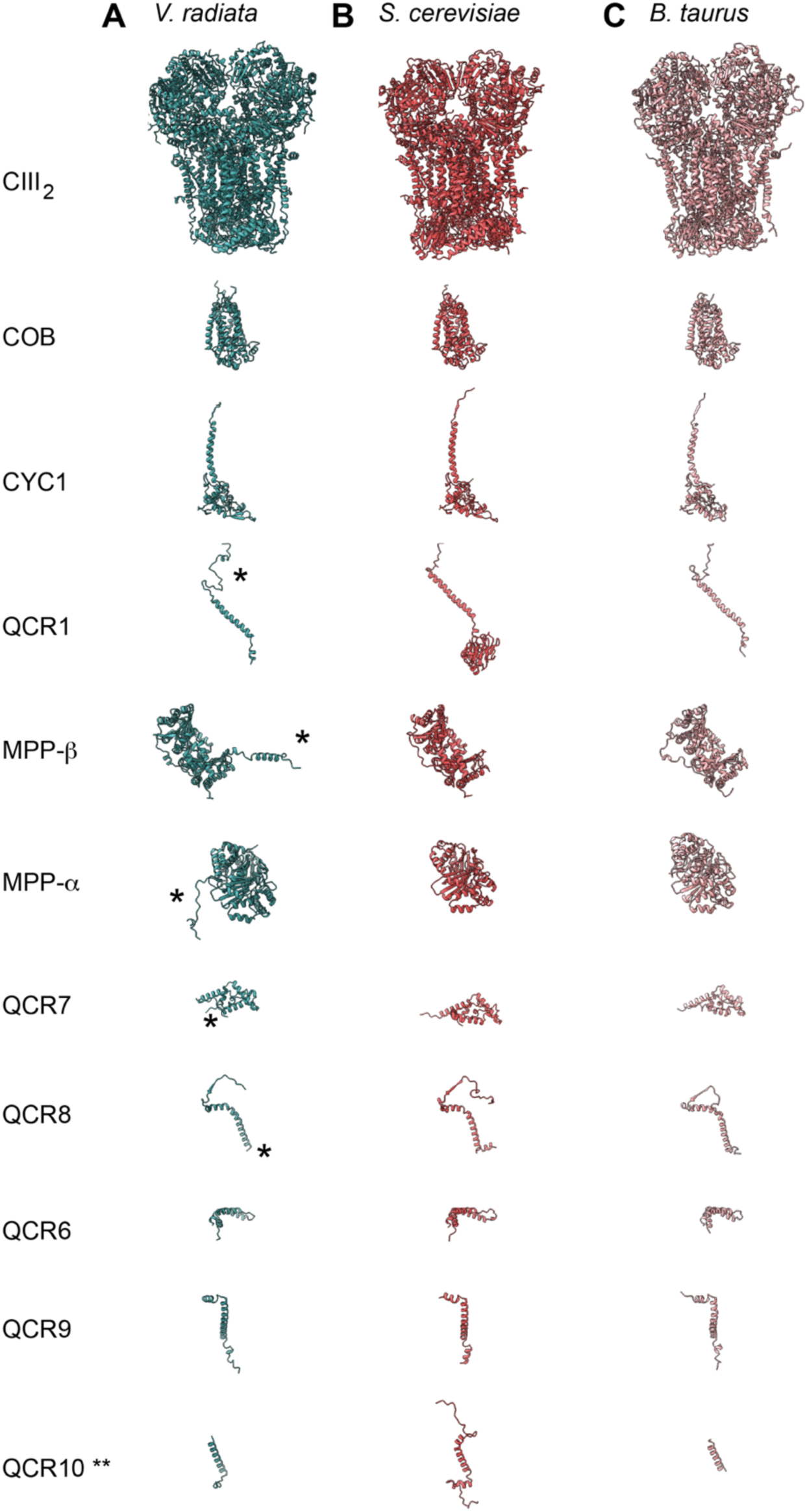
CIII subunit comparison. (A-C) CIII subunits are shown for (A) *V. radiata* (this work), (B) *S. cerevisiae* (PDB: 6HU9) and (C) *B. taurus* (PDB: 1BGY). Yeast and bovine subunits were independently aligned to the corresponding *V. radiata* subunit. The orientation of each subunit corresponds to the orientation on CIII_2_ in the top row. Note that only the atomically modelled residues are shown. Hence, the Rieske head domain of *V. radiata* and *B. taurus*, as well as the N- and C-termini of *V. radiata* QCR10 are missing (labelled with **). Differences discussed in the text are labelled with an asterisk (*).

**Figure 2-Figure Supplement 2.**
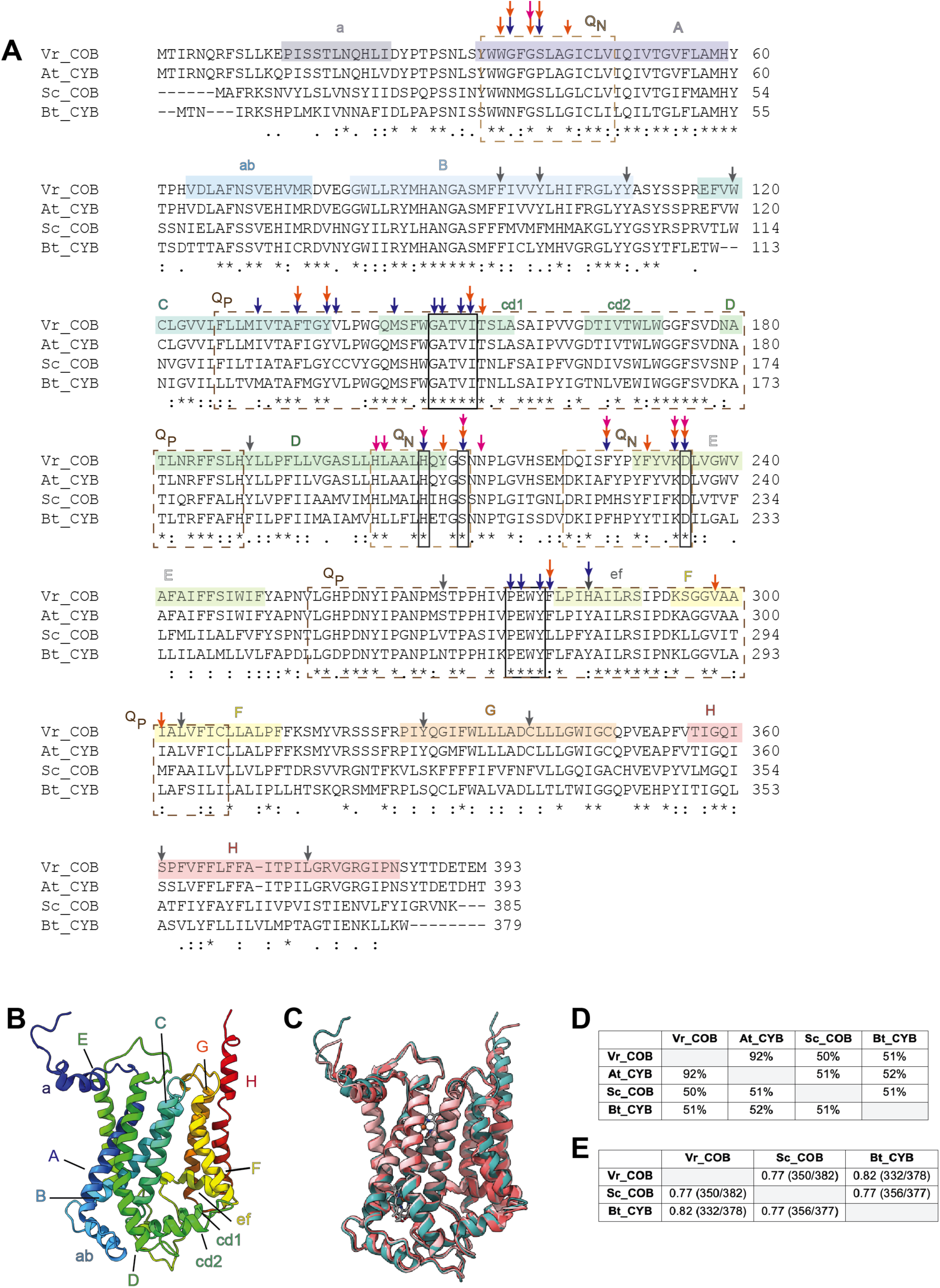
Comparison of the cyt *b* core subunit (COB) in *V. radiata* (Vr), *S. cerevisiae* (Sc), *Arabidopsis thaliana* (At) and *B. taurus* (Bt). (A) Sequence alignment of COB, performed in Clustal Omega. Secondary structure elements are highlighted in a rainbow pattern (starting with dark blue in the N-terminus to red in the C-terminus) and labelled a-F. The general locations of the residues that compose the Q_N_ site (dashed tan lines) and Q_P_ site (dashed brown lines) are labeled. Residues that directly hydrogen-bond with ubiquinone in the bovine Q_N_ site, as well as the key residues in the cd1 helix and the PEWY motif, are marked in boxes (solid lines). Other key residues, as per their role in CIII_2_ of other organisms^[15, 48, 49]^, are marked with arrows: blue for residues that establish H-bonds with inhibitors; orange for residues whose mutation confers resistance to inhibitors; magenta for residues whose mutation affects COB assembly or the re-oxidation of *b*_h_ heme. *V. radiata* residues edited in our atomic model are marked with gray arrows. See also Table 6 for further detail on the RNA edits. Symbols underneath aligned residues: `*` fully conserved, : conservation between group of strongly similar properties,. conservation between group of weakly similar properties. (B) VrCOB shown in cartoon representation, colored in rainbow pattern with the secondary structure elements labeled. (C) Superposition of COB subunits from *V. radiata* (teal), *S. cerevisiae* (dark pink, PDB 6HU9), *B. taurus* (light pink, PDB 1BGY) shown in cartoon representation. (D) Sequence identity percentages between COB subunits of the above organisms, calculated with the Clustal Omega alignment tool in Geneious. (E) Root mean square difference (RMSD) between pruned COB subunits of the above organisms, calculated in ChimeraX and shown in Å. Parentheses indicate the number of pruned atom pairs considered over the total number of atom pairs.

**Figure 2-Figure Supplement 3.**
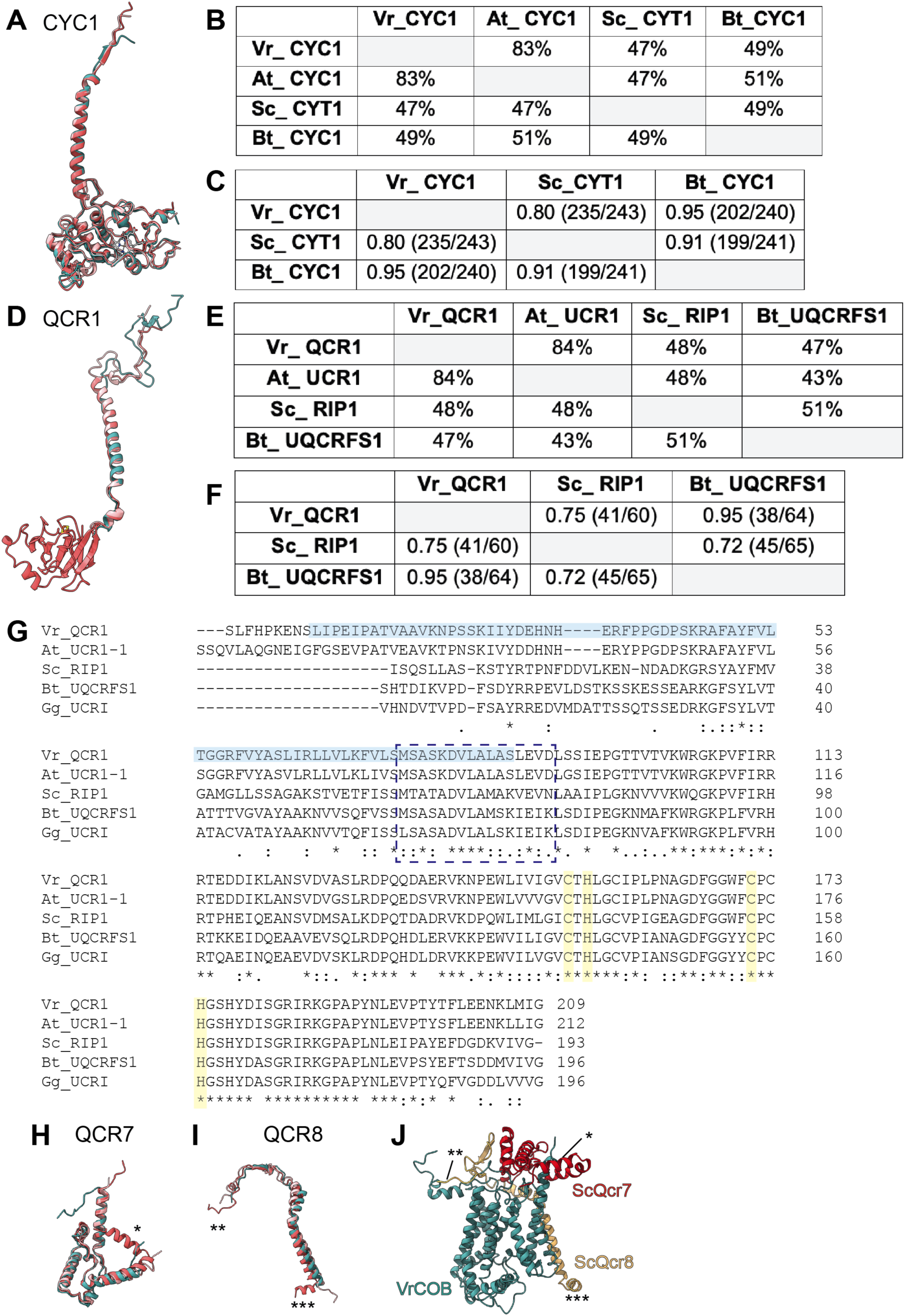
Comparison of select core and accessory subunits of CIII_2_ of *V. radiata* (Vr), *S. cerevisiae* (Sc), *A. thaliana* (At) and *B. taurus* (Bt). (A,D) Superposition of cyt *c*_1_ (CYC1) (A) and Rieske iron-sulfur protein (QCR1) (D) subunits from *V. radiata* (teal), *S. cerevisiae* (dark pink, PDB 6HU9), *B. taurus* (light pink, PDB 1BGY). (B,E) Sequence identity percentages between CYC1 (B) and QCR1 (E) subunits of the above organisms, calculated with the Clustal Omega alignment tool in Geneious. Mitochondrial import pre-sequences were not taken into account for the calculation. (C,F) Root mean square difference (RMSD) between pruned CYC1 (C) and QCR1 (F) subunits of the above organisms, calculated in ChimeraX and shown in Å. Parentheses indicate the number of pruned atom pairs considered over the total number of atom pairs. Note that the RMSD calculations in (F) do not include the head domain of VrQCR1 or Bt_UQCRFS1, as these structures do not contain atomic models for this domain. (G) Sequence alignment of QCR1 subunit (without cleaved pre-sequences) of the above organisms, with the addition of *Gallus gallus* (Gg), performed with Clustal Omega. Solid light blue box highlights the *V. radiata* residues in our atomic model. Dashed blue rectangle indicates the hinge region. The rest of the sequence corresponds to the head domain. Residues that coordinate the FeS cluster are highlighted in yellow. Symbols underneath aligned residues: `*` fully conserved, : conservation between group of strongly similar properties,. conservation between group of weakly similar properties. (H-I) Superposition of QCR7 (H) and QCR8 (I) from *V. radiata* (teal), *S. cerevisiae* (dark pink; PDB: 6HU9) and *B. taurus* (light pink; PDB: 5B1A). Subunits were aligned to the corresponding *Vigna radiata* subunit. Subunit names correspond to *V. radiata* (see also Table 4). Asterisks (*) mark the structural features present in ScQcr7 and ScQcr8 that are missing in the *V. radiata* and *B. taurus*, as discussed in the text. (J) Superposition of VrCOB (teal) with QCR7 (red) and QCR8 (yellow) from *S. cerevisiae*, aligned by the COB subunits. Asterisks correspond to the features in (I).

**Figure 3-Figure Supplement 1.**
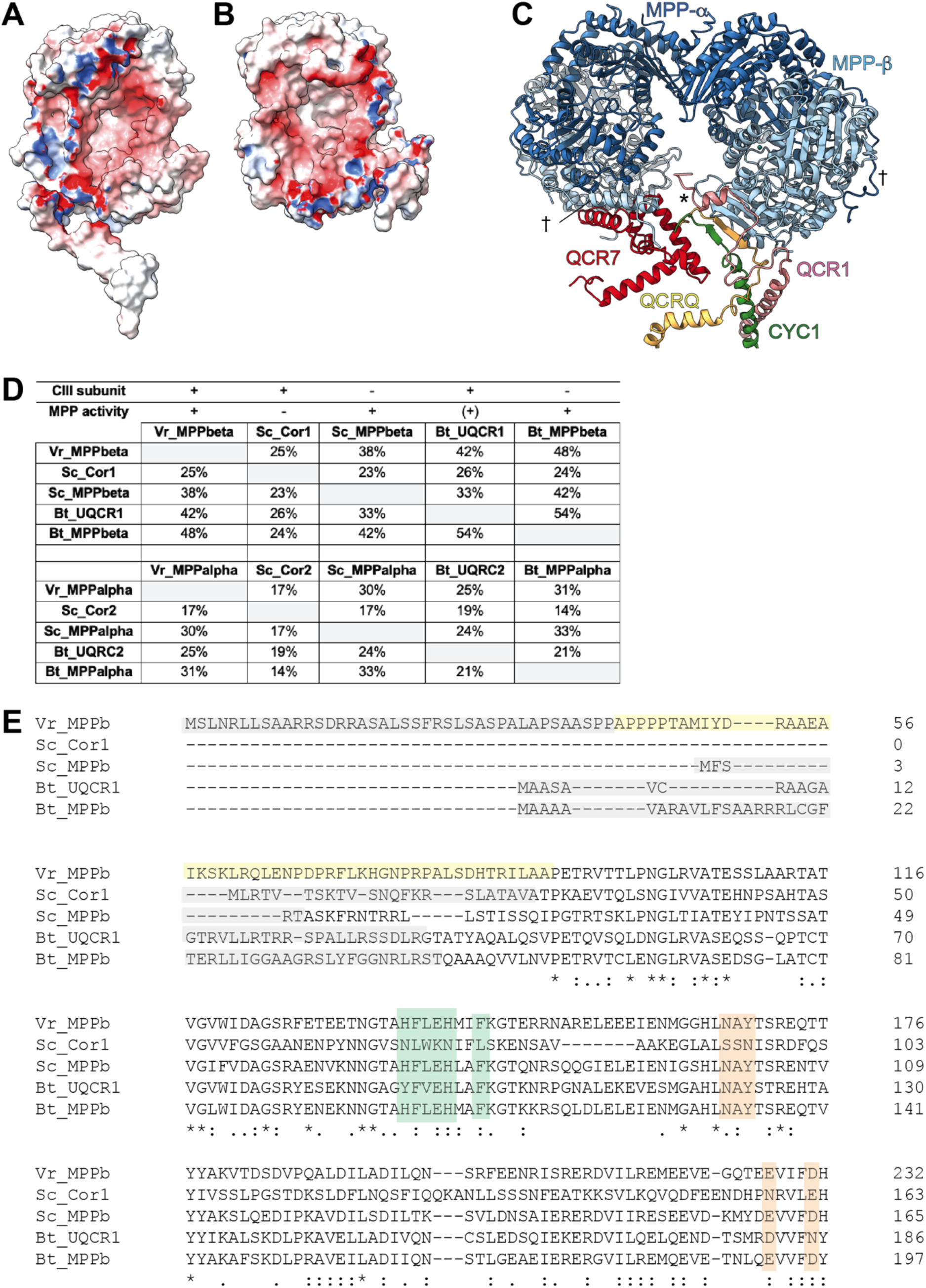
Further characterization of VrMPP subunits and their homologues. (A-B) Electrostatic potential of surface of VrMPP-*β* (A) and VrMPP-*α* (B). Electrostatic potential was calculated using Delphi^[90]^,^[91]^, with standard parameters. Red, negative; white, neutral; blue, positive. (C) VrMPP-*β*-anchoring interactions. The main anchor is marked with an asterisk (*) and comprises the *β*-sheet with strands from VrMPP-*β*, VrCYC1 and VrQCRC, as well as further interactions with the VrUCR1 N-terminus and helix from VrQCR7. The plant-specific interactions joining one MPP*α*/*β*dimer to the other (N-termini of Vr MPP-*α*and VrMPP-*β*) are marked with a cross (^†^). (D) Sequence identity percentages between MPP-*β* and MPP-*α* homologues both CIII_2_ subunits and soluble MPP in *V. radiata* (Vr), *S. cerevisiae* (Sc) and *B. taurus* (Bt). MPP subunits that are part of CIII and that show MPP enzymatic activity are marked with a plus symbol. A plus symbol in parenthesis indicates that Bt_UQCR1/2 only have known enzymatic MPP activity towards one substrate. Sequence identity percentages were calculated using the Clustal Omega multiple alignment tool in Geneious. (E) Protein sequence alignment of VrMPP-*β* subunit (VrMPPb) and its CIII_2_ and soluble MPP homologues from (D) were aligned with Clustal Omega. For space, only the first ∼200 residues of each sequence are shown. Key residues and sequences are highlighted as follows. Green: catalytic residues; orange: substrate-recognition residues; grey: cleaved import sequence (as per Uniprot entry and/or existing structure); yellow: VrMPP-*β*’s N terminal extension. Symbols underneath aligned residues: `*` fully conserved, : conservation between group of strongly similar properties,. conservation between group of weakly similar properties.

**Figure 3-Figure Supplement 2.**
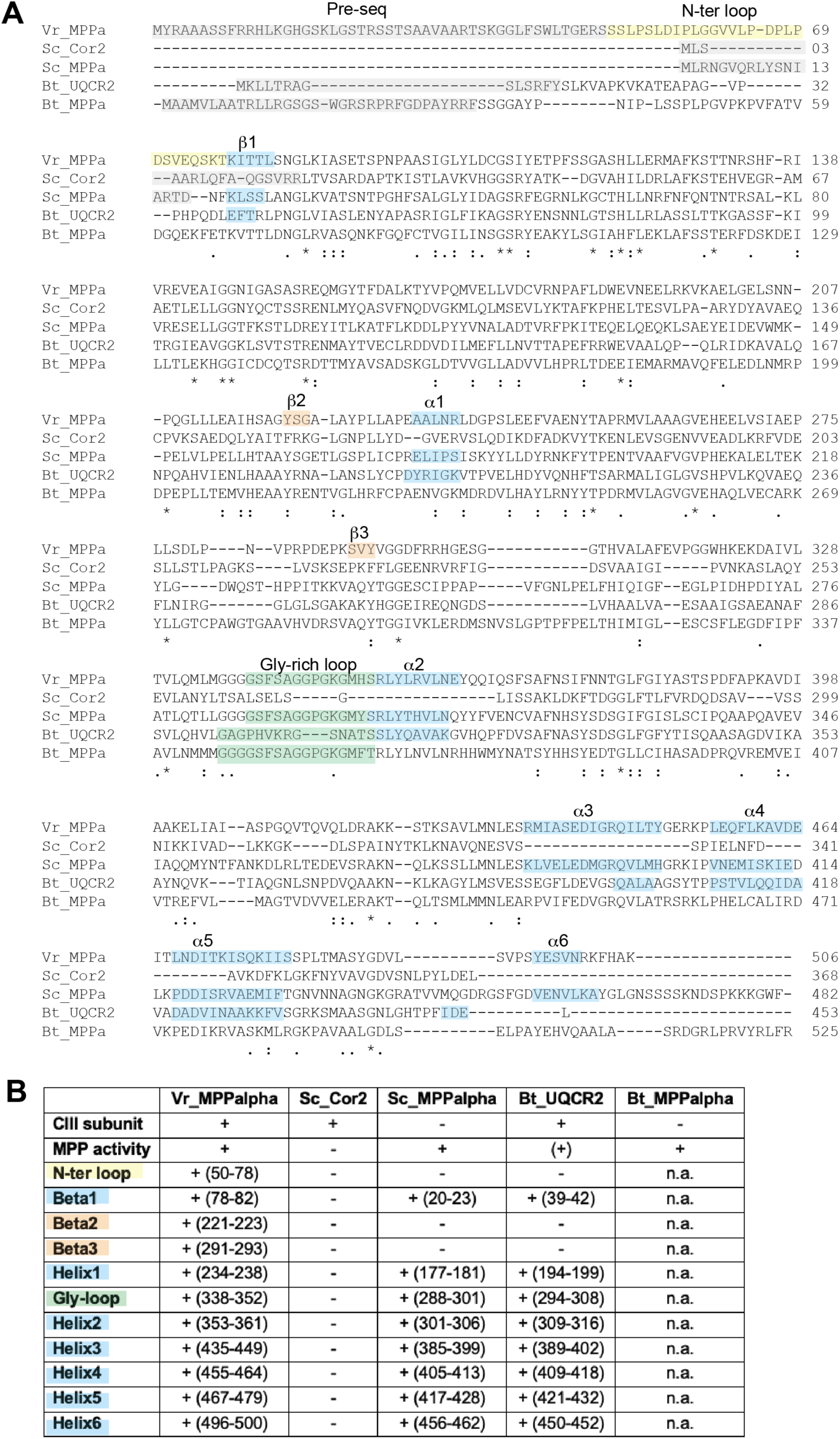
Alignment of MPP-*α* homologues. (A) Protein sequences of *V. radiata*‘s MPP-*α*subunit (VrMPPa), *S. cerevisiae* (Sc) and *B. taurus* (Bt) CIII_2_ (Cor2, UQCR2) and soluble MPP (MPPa) subunits were aligned with Clustal Omega. Key residues and sequences are highlighted as follows: grey for cleaved import pre-sequence (as per Uniprot entry and/or existing structure); yellow for VrMPP-*α*’s N terminal extension; green for substrate binding/release glycine-rich loop; blue for elements missing in ScCor2; orange for VrMPP-*α*’s plant-specific additional secondary structure elements. Symbols underneath aligned residues: `*` fully conserved, : conservation between group of strongly similar properties,. conservation between group of weakly similar properties. (B) Summary of differences in structural features of VrMPP-*α*and its homologues from (A), with the same color coding. Plus symbols (+) indicate the presence of the structural element (residue numbers in parenthesis), obtained from inspection of the structures. PDB codes of structures used: 1HR6 (ScMPP-*α*), 6HU9 (ScCor2), 1BGY (BtUQCR2). N.a. indicates that the BtMPP-*α* structure is not available and thus the comparison could not be made.

**Figure 5- Figure Supplement 1.**
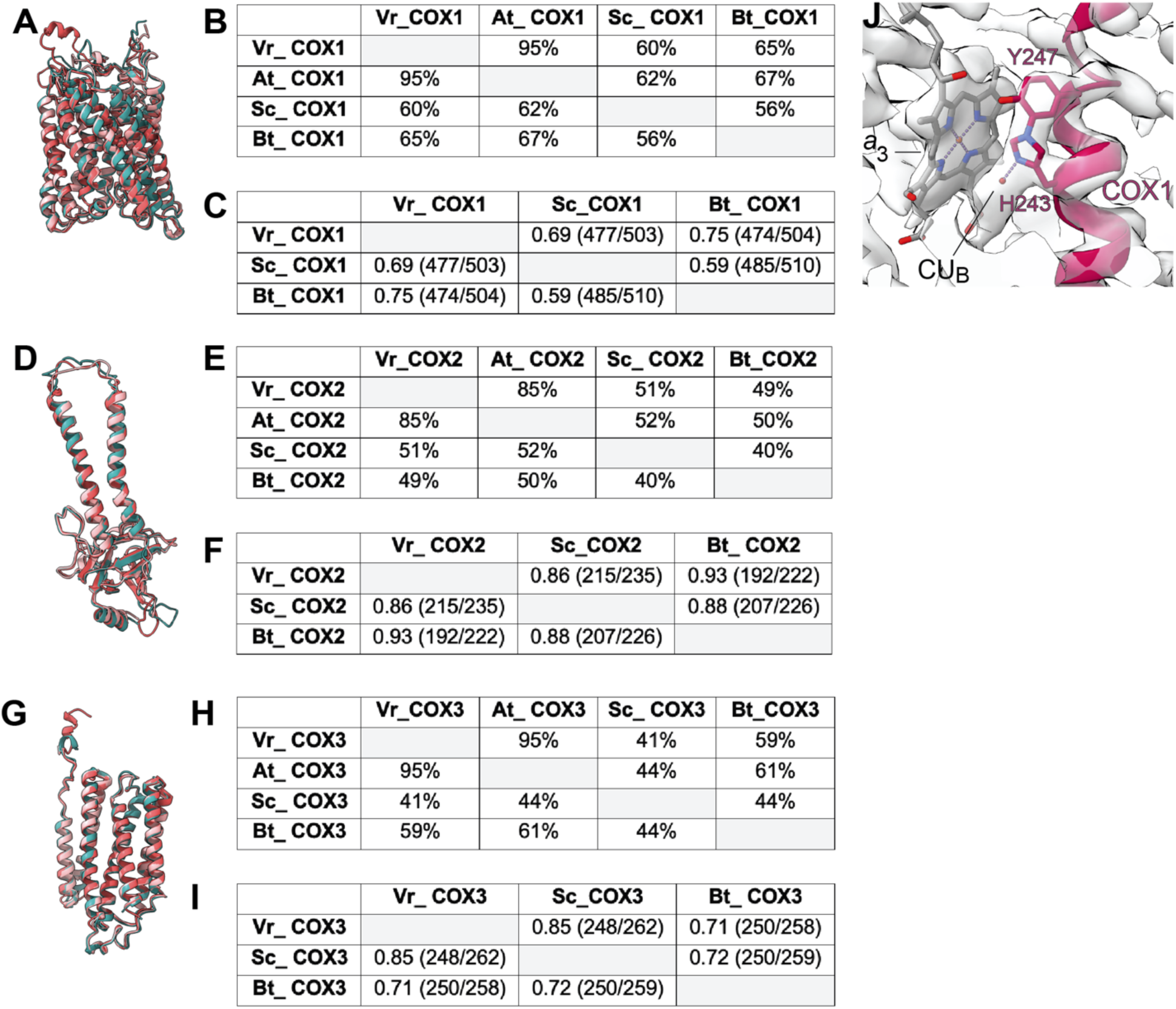
Comparison of the COX1-3 subunits in *V. radiata* (Vr), *S. cerevisiae* (Sc), *A. thaliana* (At) and *B. taurus* (Bt). (A,D,G) Superposition of COX1 (A), COX2 (D) and COX3 (G) subunits from *V. radiata* (teal), *S. cerevisiae* (dark pink, PDB 6HU9), *B. taurus* (light pink, PDB 5B1A). (B,E,H) Sequence identity percentages between COX1 (B), COX2 (E) and COX3 (H) subunits of the above organisms, calculated with the Clustal Omega alignment tool in Geneious software. (C,F,I) Root mean square difference (RMSD) between pruned COX1 (C), COX2 (F) and COX3 (I) subunits of the above organisms, calculated in ChimeraX and shown in Å. Parentheses indicate the number of pruned atom pairs considered over the total number of atom pairs. (J) HPEVY ring of VrCOX1. VrCOX1 helix shown in cartoon representation, with the covalent ring between Nε-His243 and Cε-Tyr247 shown in stick. Density shown as transparent surface. Heme a_3_ (a_3_) and CU_B_ of the dinuclear center shown as stick.

**Figure 6- Figure Supplement 1.**
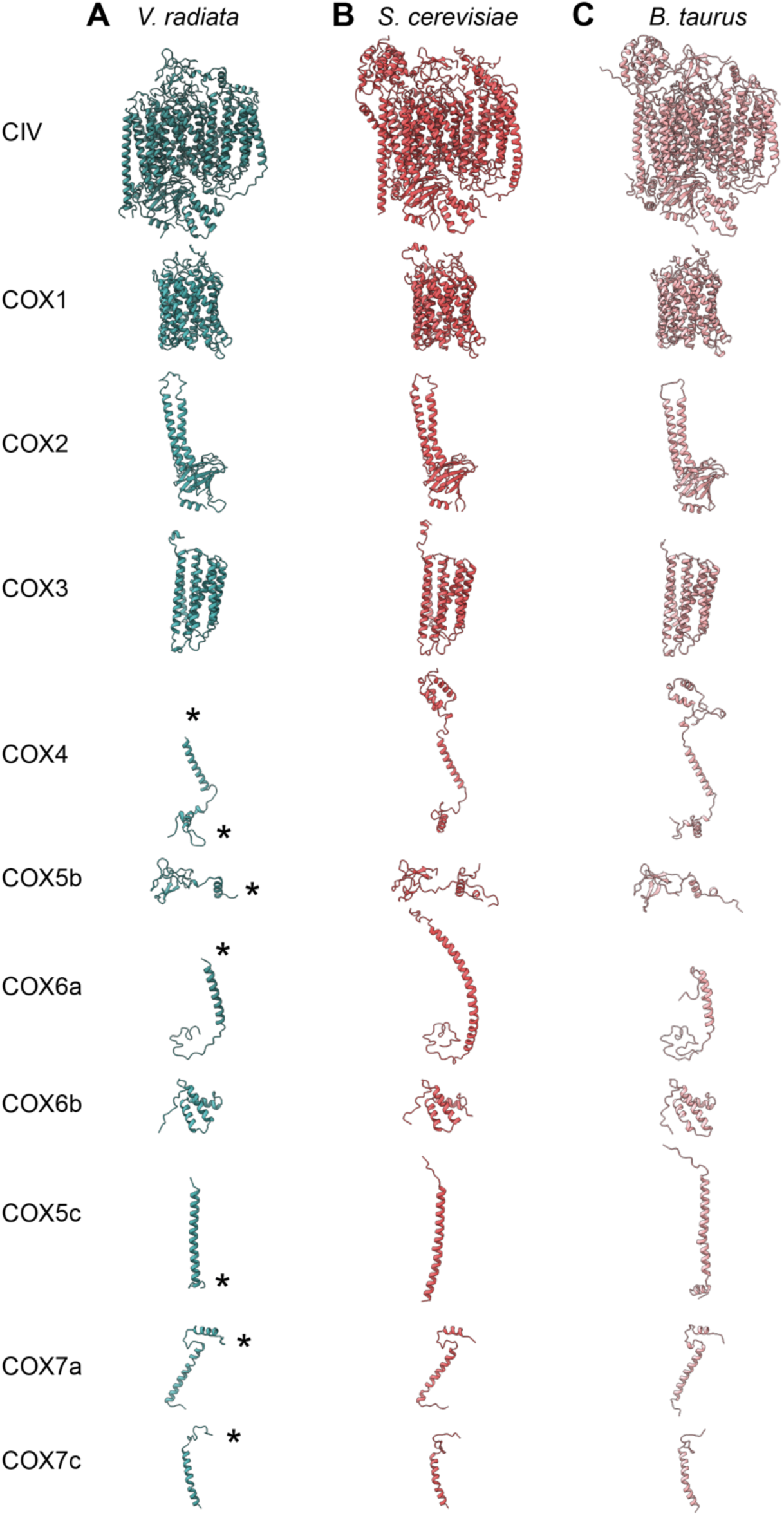
CIV subunit comparison. (A-C) CIV subunits are shown for (A) *V. radiata* (this work), (B) *S. ceresiviae* (PDB: 6HU9) and (C) *B. taurus* (PDB: 5B1A). Yeast and bovine subunits were independently aligned to the corresponding *V. radiata* subunit. The orientation of each subunit corresponds to the orientation on CIV in the top row. Only the modelled residues are shown. Differences discussed in the text are labelled with an asterisk (*).

**Figure 7- Figure Supplement 1.**
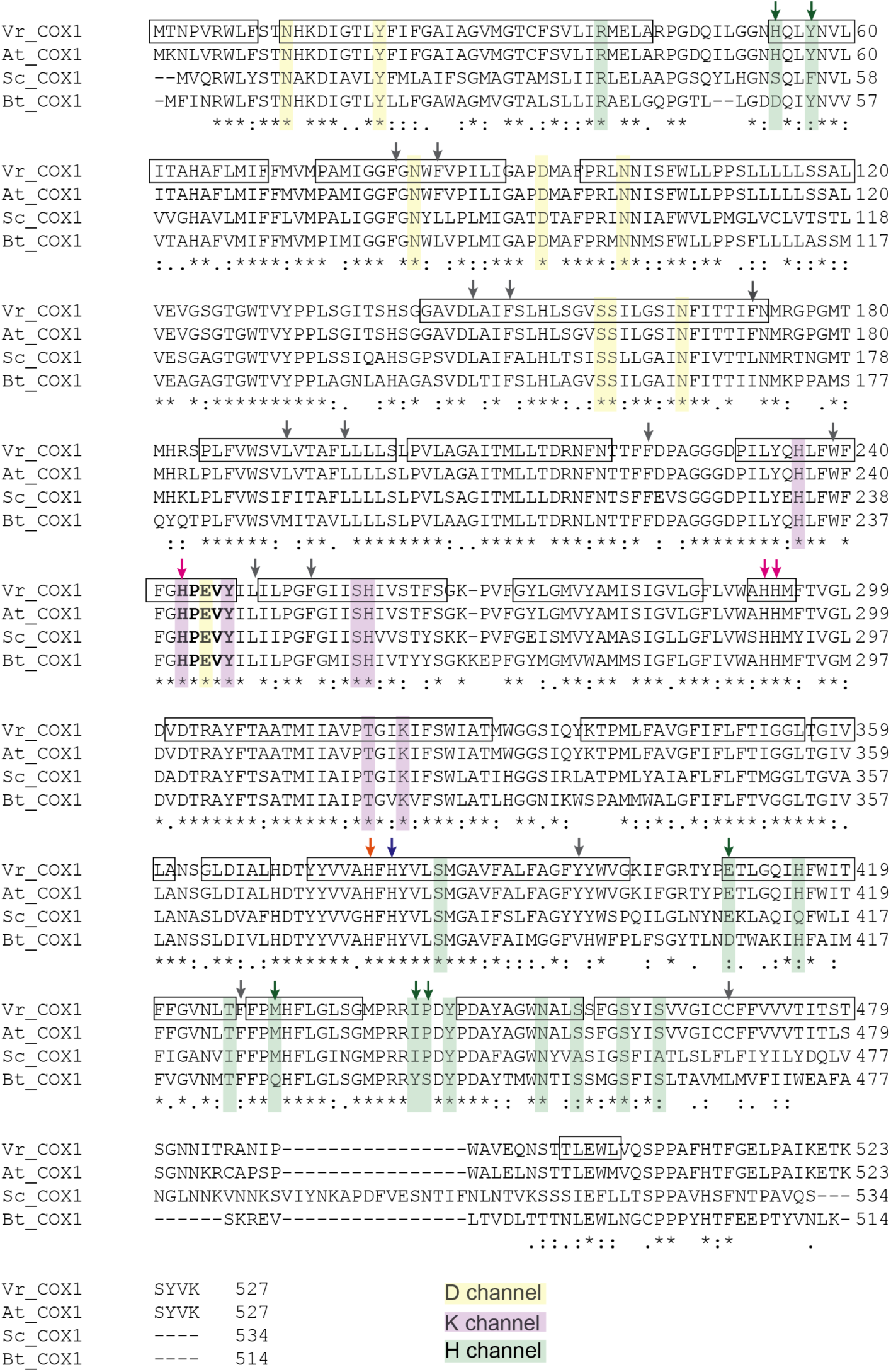
Sequence alignment of COX1 highlighting the H, D and K channels. COX1 sequences from *V. radiata* (Vr), *Arabidopsis thaliana* (At), *S. cerevisiae* (Sc), and *B. taurus* (Bt) were aligned with Clustal Omega. Proton pathway residues are highlighted: D channel in yellow, H channel in green and K channel in purple. Proton-channel residues that differ between *V. radiata* and *B. taurus* are marked with an arrow of the same color as the channel to which the residue belongs. Boxes indicate *α*-helices in the VrCOX1 atomic model. Key residues are marked with arrows: heme *a*-coordinating residues in blue, heme *a*_3_-coordinating residues in orange, CU_B_-coordinating residues in magenta. RNA editing sites are marked with gray arrows, see also Table 6. The HPEVY ring is bolded. Symbols underneath aligned residues: `*` fully conserved, : conservation between group of strongly similar properties,. conservation between group of weakly similar properties.

**Figure 8- Figure Supplement 1.**
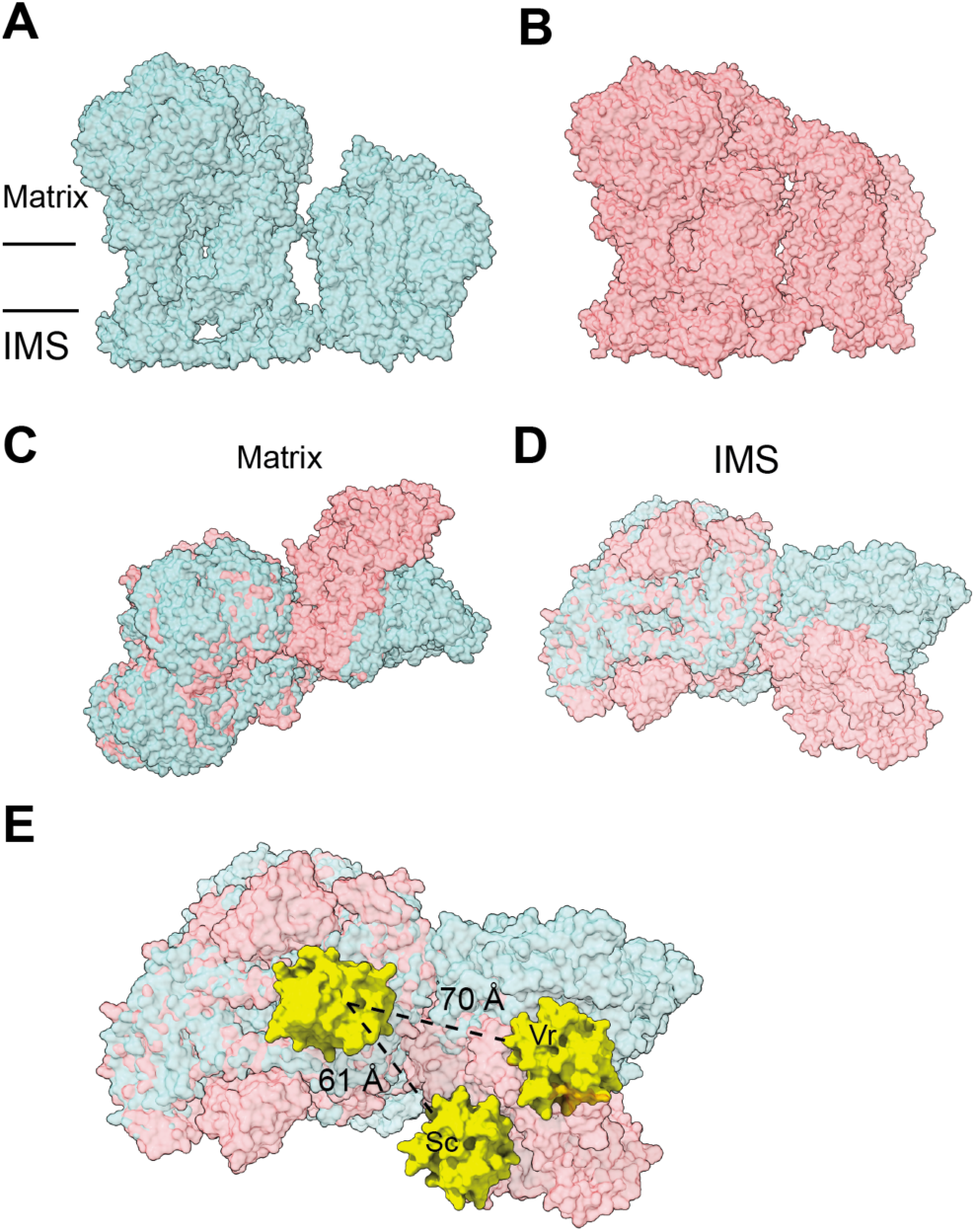
Differences in SC III_2_+IV interactions between *V. radiata* and *S. cerevisiae* (PDB: 6HU9). The supercomplexes are aligned by *V. radiata*’s COB and CYC1 and shown in surface representation. (A-B) *V. radiata* (A) and *S. cerevisiae* (B) SC III_2_+IV viewed from the membrane. (C-D) Superposed *V. radiata* (green) and *S. cerevisiae* (pink) viewed from the matrix (C) or the intermembrane space, IMS (D). (E) Cyt *c* (yellow) docked onto CIII_2_ and CIV in superposed *V. radiata* (green) or *S. cerevisiae* (pink) SC III_2_+IV viewed from the IMS. Distance between the CIII_2_- and CIV-bound cyt *c* in the *V. radiata* (Vr) and *S. cerevisiae* (Sc) supercomplexes shown. Cyt *c* was docked based on the *S. cerevisiae* CIII_2_-bound cyt *c* (1KYO) and the bovine CIV-bound cyt *c* (5IY5). CIII_2_-bound cyt c was docked onto *V. radiata* CIII_2_ by aligning CYC1. CIV-bound cyt *c* was docked onto *V. radiata* and *S. cerevisiae* CIV by aligning COX2. Distance was measured edge-to-edge of the heme conjugated ring systems.

## Materials and methods

### Key Resources Table

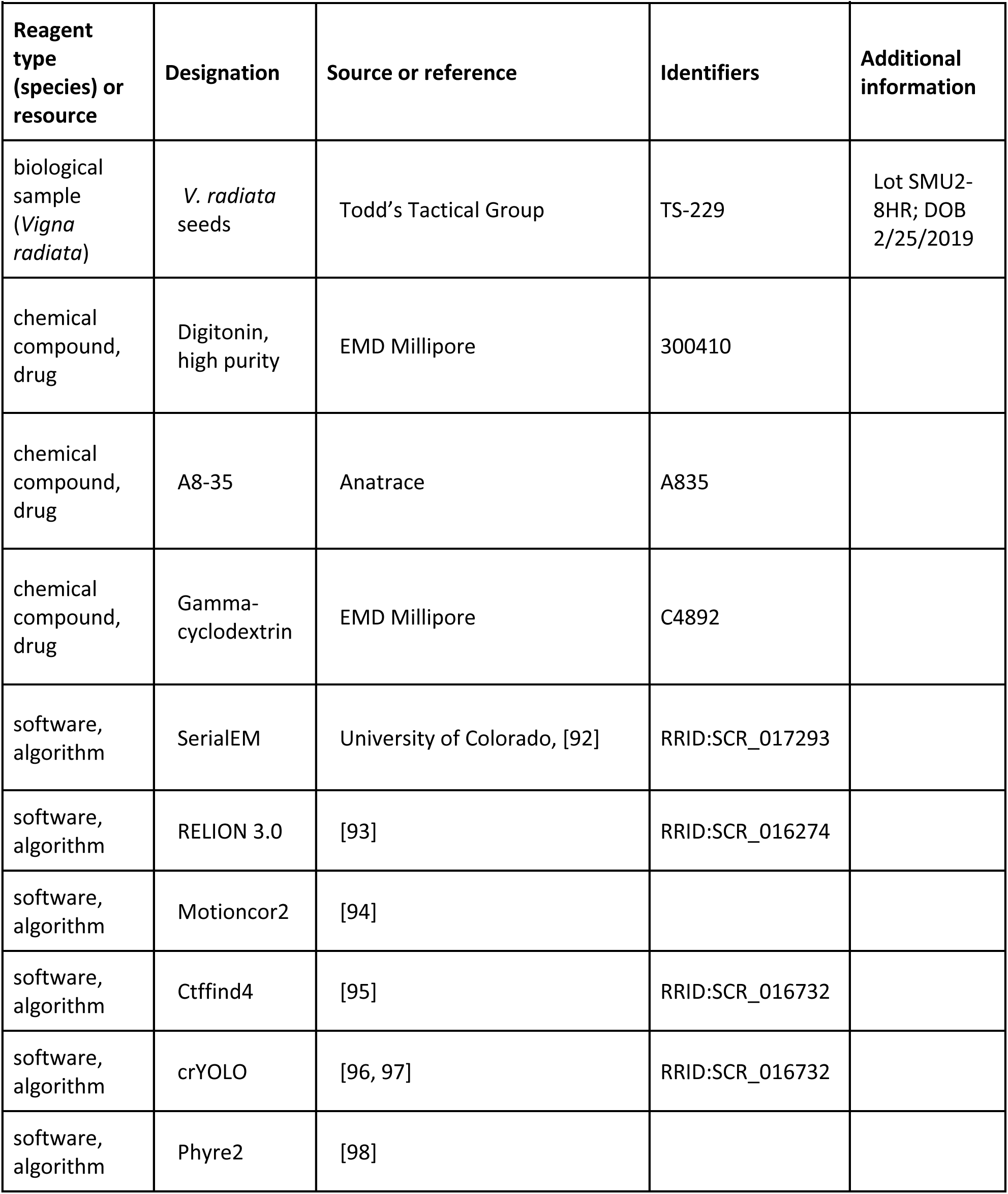
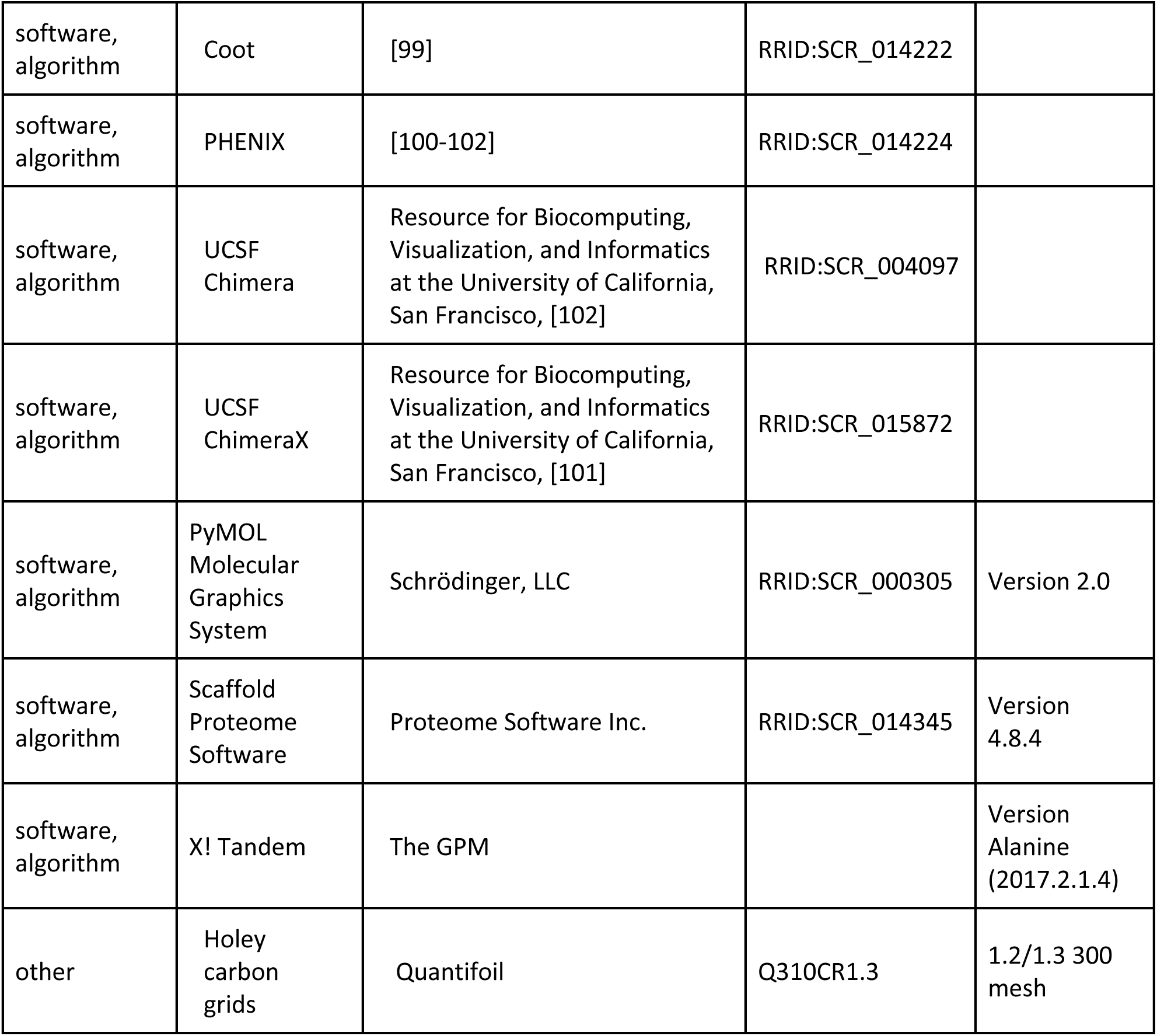

### Overall sample and data collection

The cryoEM dataset used in this paper is the same sample, grid and micrographs as those used in ref. ^[46]^. CryoEM data processing for the structures reported in this paper and those reported in ref. ^[46]^ diverged after 2D classification. Further data processing for the structures shown here is described in detail below.

### *Vigna radiata* mitochondria purification

*V. radiata* seeds were purchased from Todd’s Tactical Group (Las Vegas, Nevada, USA). Seeds were incubated in 1% (v:v) bleach for 20 minutes and rinsed until the water achieved neutral pH. Seeds were subsequently imbibed in a 6 mM CaCl_2_ solution for 20 hours in the dark. The following day, the imbibed seeds were sown in plastic trays on damp cheesecloth layers, at a density of 0.1 g/cm^2^ and incubated in the dark at 20°C for 6 days. The resulting etiolated mung beans were manually picked, and the hypocotyls were separated from the roots and cotyledons. The hypocotyls were further processed for mitochondria purification based on established protocols^[103]^. Briefly, hypocotyls were homogenized in a Waring blender with homogenization buffer (0.4 M sucrose, 1 mM EDTA, 25 mM MOPS-KOH, 10 mM tricine, 1% w:v PVP-40, freshly added 8 mM cysteine and 0.1% w:v BSA, pH 7.8) before a centrifugation of 10 min at 1,000 x *g* (4 °C). The supernatant was collected and centrifuged for 30 min at 12,000 x *g* (4 °C). The resulting pellet was resuspended with wash buffer (0.4 M sucrose, 1 mM EDTA, 25 mM MOPS-KOH, freshly added 0.1% w:v BSA, pH 7.2) and gently centrifuged at 1,000 x *g* for 5 min (4 °C). This supernatant was then centrifuged for 45 min at 12,000 x *g*. The resulting pellet was resuspended in wash buffer, loaded on to sucrose step gradients (35% w:v, 55% w:v, 75% w:v) and centrifuged for 60 min at 52,900 x *g*. The sucrose gradients were fractionated with a BioComp Piston Gradient Fractionator connected to a Gilson F203B fraction collector, following absorbance at 280 nm. The fractions containing mitochondria were pooled, diluted 1:5 in 10 mM MOPS-KOH, 1 mM EDTA, pH 7.2 and centrifuged for 20 min at 12,000 x *g* (4 °C). The pellet was resuspended in final resuspension buffer (20 mM HEPES, 50 mM NaCl, 1 mM EDTA, 10% glycerol, pH 7.5) and centrifuged for 20 min at 16,000 x *g* (4 °C). The supernatant was removed, and the pellets were frozen and stored in a −80°C freezer. The yield of these mitochondrial pellets was 0.8 –1 mg per gram of hypocotyl.

### *Vigna radiata* mitochondrial membrane wash

Frozen *V. radiata* mitochondrial pellets were thawed at 4 °C, resuspended in 10 mL of chilled (4 °C) double-distilled water per gram of pellet and homogenized with a cold Dounce glass homogenizer on ice. Chilled KCl was added to the homogenate to a final concentration of 0.15 M and further homogenized. The homogenate was centrifuged for 45 min at 32,000 x *g* (4 °C). The pellets were resuspended in cold Buffer M (20 mM Tris, 50 mM NaCl, 1 mM EDTA, 2 mM DTT, 0.002% PMSF, 10% glycerol, pH 7.4) and further homogenized before centrifugation at 32,000 x *g* for 45 min (4 °C). The pellets were resuspended in 3 mL of Buffer M per gram of starting material and further homogenized. The protein concentration of the homogenate was determined using a Pierce BCA assay kit (Thermo Fisher, Waltham, Massachusetts, USA), and the concentration was adjusted to a final concentration of 10 mg/mL and 30% glycerol.

### Extraction and purification of mitochondrial complexes

Washed membranes were thawed at 4 °C. Digitonin (EMD Millipore, Burlington, Massachusetts, USA) was added to the membranes at a final concentration of 1% (w:v) and a digitonin:protein ratio of 4:1 (w:w). Membrane complexes were extracted by tumbling this mixture for 60 min at 4 °C. The extract was centrifuged at 16,000 x *g* for 45 min (4 °C). Amphipol A8-35 (Anatrace, Maumee, Ohio, USA) was added to the supernatant at a final concentration of 0.2% w:v and tumbled for 30 min at 4°C, after which gamma-cyclodextrin (EMD Millipore, Burlington, Massachusetts, USA) was added stepwise to a final amount of 1.2x gamma-cyclodextrain:digitonin (mole:mole). The mixture was centrifuged at 137,000 x *g* for 60 min (4 °C). The supernatant was concentrated with centrifugal protein concentrators (Pall Corporation, NY, NY, USA) of 100,000 MW cut-off, loaded onto 10%-45% (w:v) or 15%-45% (w:v) linear sucrose gradients in 15 mM HEPES, 20 mM KCl, pH 7.8 produced using factory settings of a BioComp Instruments (Fredericton, Canada) gradient maker and centrifuged for 16 h at 37,000 x *g* (4 °C). The gradients were subsequently fractionated with a BioComp Piston Gradient Fractionator connected to a Gilson F203B fraction collector, following absorbance at 280 nm. For grid preparation, the relevant fractions were buffer-exchanged into 20 mM HEPES, 150 mM NaCl, 1 mM EDTA, pH 7.8 (no sucrose) and concentrated to a final protein concentration of 6 mg/mL and mixed one-to-one with the same buffer containing 0.2% digitonin (w:v), for a final concentration of 0.1% digitonin (w:v).

### Mass spectrometry

Samples were digested with the S-Trap micro (PROTIFI) digestion. Digestion followed the S-trap protocol; the proteins were reduced and alkylated, the buffer concentrations were adjusted to a final concentration of 5% SDS, 50 mM TEAB,12% phosphoric acid was added at a 1:10 (v:v) ratio with a final concentration of 1.2% and S-trap buffer (100 mM TEAB in 90% MEOH) was added at a 1:7 ratio (v:v) ratio. The protein lysate S-trap buffer mixture was then spun through the S-trap column and washed 3 times with S-Trap buffer. Finally, 50 mM TEAB and 1µg of trypsin (1:25 ratio) was added and the sample is incubated overnight with one addition of 50 mM TEAB and trypsin after two hours. The following day the digested peptides were released from the S-trap solid support by spinning at 1 minute for 3,000 x g with a series of solutions starting with 50 mM TEAB which is placed on top of the digestion solution then 5% formic acid followed by 50% acetonitrile, 0.1% formic acid. The solution is then vacuum centrifuged to almost dryness and resuspended in 2% acetonitrile, 0.1% triflouroacetic acid (TFA) and subjected to Fluorescent Peptide Quantification (Pierce).

Digested peptides were analyzed by LC-MS/MS on a Thermo Scientific Q Exactive plus Orbitrap Mass spectrometer in conjunction Proxeon Easy-nLC II HPLC (Thermo Scientific) and Proxeon nanospray source. The digested peptides were loaded on a 100 micron x 25 mm Dr. Masic reverse phase trap where they were desalted online before being separated using a 75 micron x 150 mm Magic C18 200 Å 3U reverse phase column. Peptides were eluted using a 70-minute gradient with a flow rate of 300 nL/min. An MS survey scan was obtained for the m/z range 300-1600, MS/MS spectra were acquired using a top 15 method, where the top 15 ions in the MS spectra were subjected to HCD (High Energy Collisional Dissociation). An isolation mass window of 2.0 m/z was for the precursor ion selection, and normalized collision energy of 27% was used for fragmentation. A twenty second duration was used for the dynamic exclusion.

Tandem mass spectra were extracted and charge state deconvoluted by Proteome Discoverer (Thermo Scientific). All MS/MS samples were analyzed using X! Tandem (The GPM, thegpm.org; version X! Tandem Alanine (2017.2.1.4)). X! Tandem was set up to search the Uniprot *Vigna radiata* database (October 2019, 35065 entries) the cRAP database of common laboratory contaminants (www.thegpm.org/crap; 117 entries) plus an equal number of reverse protein sequences assuming the digestion enzyme trypsin. X! Tandem was searched with a fragment ion mass tolerance of 20 PPM and a parent ion tolerance of 20 PPM. Carbamidomethyl of cysteine and selenocysteine was specified in X! Tandem as a fixed modification. Glu->pyro-Glu of the n-terminus, ammonia-loss of the n-terminus, gln->pyro-Glu of the n-terminus, deamidated of asparagine and glutamine, oxidation of methionine and tryptophan and dioxidation of methionine and tryptophan were specified in X! Tandem as variable modifications.

Scaffold (version Scaffold_4.8.4, Proteome Software Inc., Portland, OR) was used to validate MS/MS based peptide and protein identifications. Peptide identifications were accepted if they could be established at greater than 98.0% probability by the Scaffold Local FDR algorithm. X! Tandem identifications required score of at least 2. Protein identifications were accepted if they could be established at greater than 6.0% probability to achieve an FDR less than 5.0% and contained at least 2 identified peptides. Protein probabilities were assigned by the Protein Prophet algorithm^[104]^. Proteins that contained similar peptides and could not be differentiated based on MS/MS analysis alone were grouped to satisfy the principles of parsimony. Proteins sharing significant peptide evidence were grouped into clusters.

### CryoEM data acquisition

The sample (6 mg/mL protein in 20 mM HEPES, 150 mM NaCl, 1 mM EDTA, 0.1% digitonin, pH 7.8) was applied onto glow-discharged holey carbon grids (Quantifoil, 1.2/1.3 300 mesh) followed by a 60 s incubation and blotting for 9 s at 15 °C with 100% humidity and flash-freezing in liquid ethane using a FEI Vitrobot Mach III.

CryoEM data acquisition was performed on a 300 kV Titan Krios electron microscope equipped with an energy filter and a K3 detector at the UCSF W.M. Keck Foundation Advanced Microscopy Laboratory, accessed through the Bay Area CryoEM Consortium. Automated data collection was performed with the SerialEM package^[92]^. Micrographs were recorded at a nominal magnification of 117,813 X, resulting in a pixel size of 0.8332 Å^2^. Defocus values varied from 1.5 to 3.0 µm. The dose rate was 20 electrons per pixel per second. Exposures of 3 s were dose-fractionated into 118 frames, leading to a dose of 0.72 electrons per Å^2^ per frame and a total accumulated dose of 51 electrons per Å^2^. A total of 9,816 micrographs were collected.

### Data processing

Software used in the project was installed and configured by SBGrid^[105]^. All processing steps were done using cryoSPARC and RELION 3.0 (ref. ^[93]^, ^[106]^) unless otherwise stated. Motioncor2 (ref. ^[94]^) was used for whole-image drift correction of each micrograph. Contrast transfer function (CTF) parameters of the corrected micrographs were estimated using Ctffind4 (ref. ^[95]^). After motion correction and CTF correction, a set of 8,541 micrographs was selected for further processing. Automated particle picking using crYOLO^[96, 97]^ resulted in ∼1.5 million particles. The particles were extracted using 400^2^ pixel box binned two-fold and sorted by reference-free 2D classification in Relion using (--max_sig 5), followed by re-extraction at 512^2^ pixel box. Reference-free 2D classification in Relion resulted in the identification of 502,224 particles that were then imported into cryoSPARC for further reference-free 2D classification. A set of 121,702 particles were identified by 2D classification in cryoSPARC to contain CIII_2_ alone or SC III_2_+IV (Figure 1-Figure Supplement 1). These particles were subjected to *ab initio* model generation with four targets to remove contaminant particles resulting in a set of 99,937 particles across three classes. Each individual class was subjected to an additional round of *ab initio* model generation with three targets. This separated CIII_2_ alone particles from the SC III2+IV class and allowed the recovery of CIII_2_ alone particles from the poor particle class. CIII_2_ alone particles from across the *ab initio* model generation jobs were pooled, defining a final class of 48,111 particles. The multiple rounds of *ab initio* model generation resulted in only one good class of 28,020 SC III_2_+IV particles. Poses for these two particle sets (CIII_2_ alone and SC III_2_+IV) were refined using cryoSPARC’s Homogeneous Refinement (New) algorithm including Defocus Refinement and Global CTF refinement. This resulted in reconstructions at 3.2 Å and 3.8 Å resolution for CIII_2_ alone and SC III2+IV respectively, according to the gold standard FSC criteria (Figure 1-Figure Supplement 1)^[107]^.

In parallel, a set of 69,876 particles were identified by further 2D and 3D classification in Relion (Figure 1-Figure Supplement 2). These particles, which contained a mixture of SC III_2_+IV and CIII_2_ alone particles, were aligned using a SC III_2_+IV model and mask. They were then subjected to five rounds of masked classification using a CIV model and mask aligned to the position of CIV in the SC. Three parallel masked classifications (all using T=8) varied in the degree of rotational searches, with either no searches, 0.1° sampling interval over +/- 0.2° search range and 3.7° sampling interval over +/- 7.5° search range. The masked 3D classification without searches was repeated successively for three rounds, inputting the best particles from the previous round into the subsequent round. The best CIV class from each 3D classification were selected and combined while removing overlaps (any particle within 200 pixels of another was considered as an overlap and discarded). This 3D classification strategy resulted in a set of 38,410 particles. The coordinates of these particles were used to extract two sets of re-centered SC particles, one centered on CIII_2_ and one centered on CIV. These two sets of particles were independently 3D-refined, CTF-refined and Bayesian-polished using a model and mask centered around CIII_2_ or CIV respectively. The CIV-centered shiny particles were subjected to a final round of 3D classification, defining a final set of 29,348 CIV particles. Although this final round of 3D classification did not improve the nominal resolution of the map significantly, it increased map quality at the periphery of the complex. These final CIII_2_ and CIV classes resulted in reconstructions at 3.7 Å and 3.8 Å resolution for CIII_2_ and IV from the SC respectively, according to the gold standard FSC criteria (Figure 1-Figure Supplement 1)^[107]^. These two maps were aligned to the full SC map and combined to make a composite map using Phenix.

3D variability analysis (3DVA) was performed on CIII_2_ alone in cryoSPARC using their built-in algorithm^[45]^. Two separate instances of 3DVA were performed, each solving for the four largest principal components. The first instance used a mask around the entire CIII_2_ and data was low-pass filtered to 6 Å resolution to remove the influence of high resolution noise from the amphipol detergent belt. The second instance used a mask focused around the IMS domain of CIII_2_ and was low-pass filtered to 5 Å resolution. All other parameters were kept as default.

### Model building and refinement

Starting template models for *V. radiata* CIII_2_ was ovine CIII_2_ (PDB: 6Q9E). Starting template models for *V. radiata* CIV were from *S. cerevisiae* (PDB: 6HU9) and bovine (PDB: 5B1A) CIV. Additionally, starting models for the *V. radiata* subunits were generated using the Phyre2 web portal^[98]^. Real-space refinement of the model was done in PHENIX^[100–102]^ and group atomic displacement parameters (ADPs) were refined in reciprocal space. The single cycle of group ADP refinement was followed by three cycles of global minimization, followed by an additional cycle of group ADP refinement and finally three cycles of global minimization^[11]^. The refined CIII_2_ and CIV models were docked into the SC III_2_+CIV map without subsequent refinement.

### Model interpretation and figure preparation

Molecular graphics and analyses were performed with UCSF Chimera^[102]^ and ChimeraX^[101]^ developed by the Resource for Biocomputing, Visualization, and Informatics at the University of California, San Francisco, with support from NIH P41-GM103311 and R01-GM129325 and the Office of Cyber Infrastructure and Computational Biology, National Institute of Allergy and Infectious Diseases. PyMOL Molecular Graphics System, Version 2.0 Schrödinger, LLC was also used.

## Acknowledgments

We are grateful to K. Abe, M.G. Zaragoza, C. Goodwin, R. Murguia and C. Bower for help with *V. radiata* growth and mitochondrial isolations. Data was collected at the UCSF W.M. Keck Foundation Advanced Microscopy Laboratory, accessed through the Bay Area CryoEM Consortium BACEM, with the assistance of D. Bulkley and Z. Yu. We are grateful to W. Broadly of the UC Davis High-Performance Cluster and M. Salemi of the UC Davis Proteomics Core for technical assistance. M.M. acknowledges funding from the UC Davis POP Program.

